# Associations between MHC class II variation and phenotypic traits in a free-living sheep population

**DOI:** 10.1101/2021.04.22.440962

**Authors:** Wei Huang, Kara L Dicks, Keith T Ballingall, Susan E Johnston, Alexandra M Sparks, Kathryn Watt, Jill G. Pilkington, Josephine M Pemberton

## Abstract

Pathogen-mediated selection (PMS) is thought to maintain the high level of allelic diversity observed in the major histocompatibility complex (MHC) class II genes. A comprehensive way to demonstrate contemporary selection is to examine associations between MHC variation and individual fitness. As individual fitness is hard to measure, many studies examine associations between MHC diversity and phenotypic traits which include direct or indirect measures of adaptive immunity thought to contribute to fitness. Here, we tested associations between MHC class II variation and five representative phenotypic traits measured in August: weight, strongyle faecal egg count, and plasma IgA, IgE and IgG immunoglobulin titres against the gastrointestinal nematode parasite *Teladorsagia circumcincta* in a free-living population of Soay sheep. We found no association between MHC class II variation and August weight or strongyle faecal egg count. We did however find associations between MHC class II variation and immunoglobulin levels which varied with age, isotype and sex. Our results suggest associations between MHC and phenotypic traits are more likely to be found for traits more closely associated with pathogen defence than integrative traits such as body weight and highlight a useful role of MHC-antibody associations in examining selection on MHC genes.

## Introduction

The immune system provides a variety of mechanisms to protect the host from infection by rapidly evolving and highly variable pathogens. The diversity of immune-related proteins and their associated genes are believed to have evolved in response to such pathogen diversity via the process of coevolution (Eizaguirre, Lenz et al. 2012, Pilosof, Fortuna et al. 2014). Among the proteins directly involved in the initiation of adaptive immunity, major histocompatibility complex (MHC) molecules, encoded by MHC class I and class II gene families, are the most variable and have been intensively researched in many species (Edwards and Hedrick 1998, Bernatchez and Landry 2003, Piertney and Oliver 2006). MHC genes encode heterodimeric MHC molecules which bind and present short peptides derived from pathogens to T cells to invoke and coordinate the adaptive immune response. Classical MHC class I genes are primarily responsible for presenting peptides derived from intracellular pathogens such as viruses, while classical MHC class II genes typically present peptides derived from extracellular pathogens such as bacteria and parasites (Bernatchez and Landry 2003).The tight mechanistic link between pathogen infection and MHC molecules leads to the expectation that selection pressure imposed by pathogens, known as pathogen-mediated selection (PMS), is the major force driving high levels of diversity at the MHC (Bernatchez and Landry 2003, Piertney and Oliver 2006, Spurgin and Richardson 2010).

Substantial effort has been made to investigate how pathogen-mediated selection (PMS) can maintain high levels of MHC diversity across a wide variety of vertebrate taxa (reviewed by (Bernatchez and Landry 2003, Piertney and Oliver 2006, Spurgin and Richardson 2010)). There are several non-mutually-exclusive mechanisms by which pathogen-mediated selection may occur. 1) Heterozygote advantage (HA): The HA occurs when heterozygotes have greater fitness than either homozygote (Hughes and Nei 1988, Takahata and Nei 1990, Penn, Damjanovich et al. 2002). 2) Divergent allele advantage (DAA): DAA is an extension of HA. Under DAA, individuals with high levels of functional divergence between MHC alleles have a selective advantage over individuals with lower levels of allelic divergence (Wakeland, Boehme et al. 1990). 3) Negative frequency-dependent selection (NFDS): NFDS occurs due to rare allele advantage; pathogens are predicted to be under selection to evade the most common MHC alleles, resulting in rare MHC alleles having a selective advantage, and creating a cyclical co-evolutionary arms race (Takahata and Nei 1990, Slade and Mccallum 1992). 4) Fluctuating selection (FS): Under FS, directional selection due to variation in pathogen pressure varies in time and space such that it maintains diversity (Hedrick 2002). Several studies have used experimental methods to examine PMS on MHC genes (Bolnick and Stutz 2017, Phillips, Cable et al. 2018). However, experimental studies are rarely capable of replicating the wide array of pathogens and parasites that occur within a wild host and are limited in the conclusions that they can draw about natural processes. Therefore, testing these hypotheses within wild systems is valuable (Piertney and Oliver 2006, Spurgin and Richardson 2010).

A direct way to demonstrate contemporary selection on MHC genes in a wild population is to examine associations between MHC heterozygosity or genotypes and fitness. As fitness measurements are not always available, we could alternatively examine associations between MHC variation and pathogen load (Spurgin and Richardson 2010) and other fitness-related phenotypic traits, e.g. body weight. However, examining phenotypic traits is a less direct approach than examining fitness components and, in the quest to understand selection mechanisms, it is of interest to know how consistent results from these two approaches are. Only a few studies have analysed both MHC-fitness associations and MHC-phenotypic trait associations in the wild (summarized in Table 1). Of these, some studies found coherent results between MHC-fitness and MHC-phenotypic trait associations (Paterson, Wilson et al. 1998, Kloch, Baran et al. 2012, Sepil, Lachish et al. 2013), while other studies did not (Dunn, Bollmer et al. 2013). These mixed results could be due to some MHC-fitness associations not acting through the phenotypic traits examined. Therefore, it is of interest to determine which types of traits MHC-based selection is most likely to be acting through.

**Table 1.**
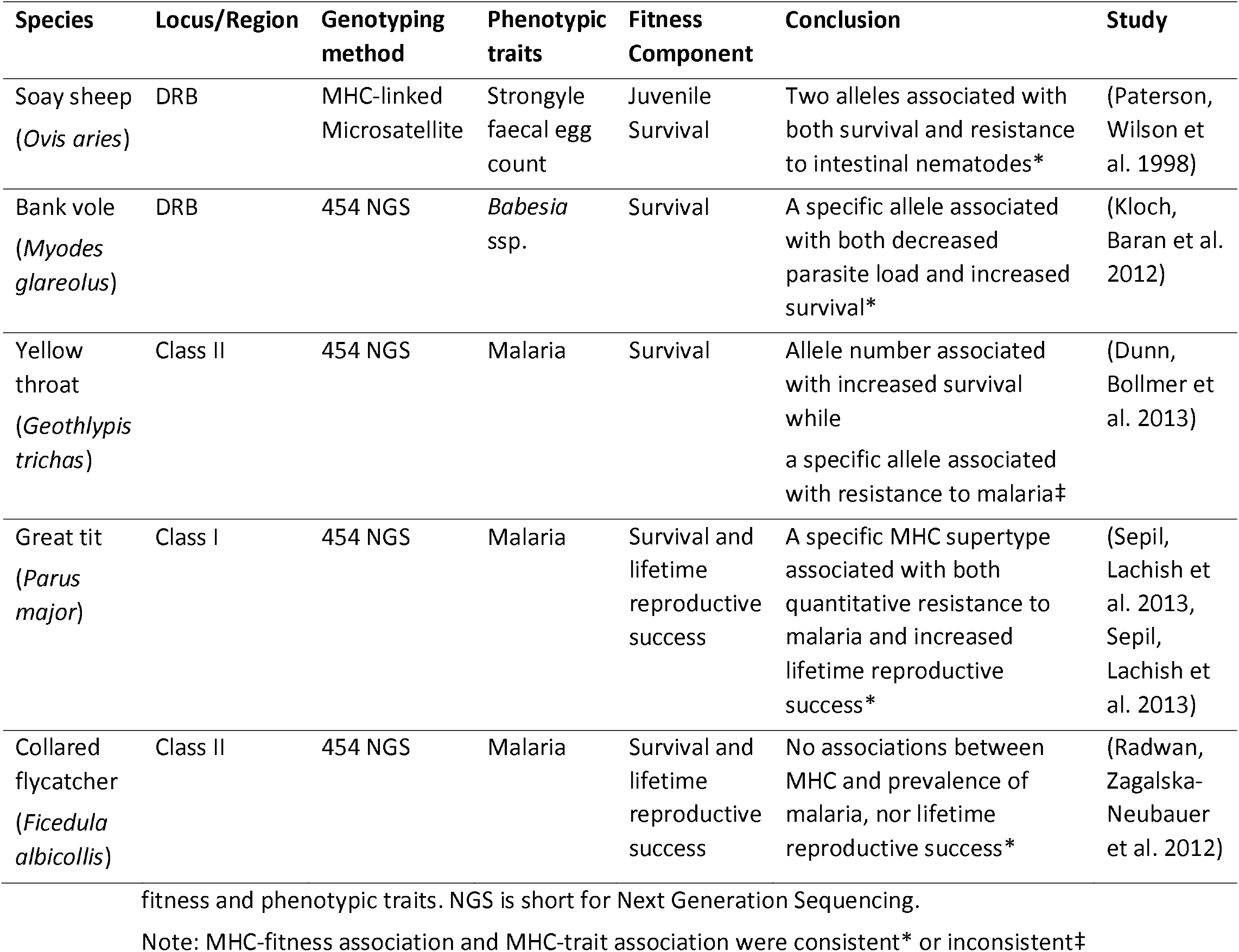
Summary of previous studies testing associations between MHC diversity and both

Another possible explanation for variation between MHC-fitness and MHC-phenotype studies may lie in analysis methods. First, MHC–phenotypic trait associations may vary due to heterogeneity in exposure or response to pathogens due to host age and sex. For example, in a recent study of black-legged kittiwake (*Rissa tridactyla*), MHC class II diversity was positively associated with growth and tick clearance in female but not in male chicks (Pineaux, Merkling et al. 2020). Therefore it is important to investigate whether MHC-phenotypic traits associations vary with age and sex. In addition, there is additive genetic variation for most phenotypic traits. When examining MHC-phenotypic trait associations in datasets including many related individuals, animal models should be used. The animal model framework includes phenotypic information from individuals of varying relatedness to estimate the additive genetic component of the trait by including the breeding value as a random effect within a mixed effect model such that variation in the trait that is due to additive genetic effects located throughout the genome could be controlled (Wilson, Reale et al. 2010). This will reduce risk of generating false positive associations MHC–trait associations.

The unmanaged Soay sheep (*Ovis aries*) population on Hirta, St Kilda, UK has been intensively studied for more than three decades (Clutton-Brock and Pemberton 2004). Since 1985, nearly all individuals living in the Village Bay study area have been followed from birth, through all breeding attempts, to death. These data, combined with a genetically-inferred multigenerational pedigree and phenotypic data for body weight, parasite load and plasma antibodies (Hayward, Wilson et al. 2011, Hayward, Nussey et al. 2014, Nussey, Watt et al. 2014, Berenos, Ellis et al. 2015, Sparks, Watt et al. 2018), enable us to investigate the interplay between MHC variation, fitness and phenotypic traits. A previous study of Soay sheep alive between 1985 and 1994 found negative associations between two alleles at an MHC-linked microsatellite and a key parasite measurement, strongyle faecal egg count (FEC), and these two alleles were also positively associated with juvenile survival (Paterson, Wilson et al. 1998). Recently, we reported associations between Soay sheep MHC class II variation and fitness measurements (Huang, Dicks et al. 2020). These new findings were enabled by major advances in the characterisation of MHC variation in Soay sheep. A total of eight MHC class II haplotypes (named A-H) were identified through sequence-based genotyping of a subset of the population (Dicks, Pemberton et al. 2019) and imputed successfully for 5349 sheep sampled from 1985 to 2012 using 13 SNPs (Dicks 2017, Dicks, Pemberton et al. 2019). We found haplotype C and D are associated with decreased and increased male total fitness (measured as the number of offspring that an individual had throughout its life span) respectively. In term of fitness components, we found MHC divergence (measured as the proportion of the amino acid sequence that differed between the two MHC haplotypes of each individual) was positively associated with juvenile survival. We also found that a haplotype (C) is associated with decreased adult male breeding success while another haplotype (F) is associated with decreased adult female life span. In addition, the frequency of haplotype D has increased significantly through the study period more than expected by drift (Huang, Dicks et al. 2020). These results indicate that there is contemporary selection on MHC class II variation in Soay sheep.

In the present study, with larger sample sizes and improved genetic resolution of haplotypes compared with the previous study (Paterson, Wilson et al. 1998), we examine the associations between MHC variation and five representative phenotypic traits in Soay sheep. These traits are August weight, a fitness-related non-immune trait, strongyle faecal egg count, FEC, a fitness-related trait with a strong link to the immune system and three immune traits, *Teladorsagia circumcincta-specific* immunoglobulin isotypes IgA, IgE and IgG (henceforth, ‘anti-*T.circ* antibodies’). August weight is an important measure of body condition in Soay sheep and high weight is advantageous for both survival and fecundity in Soay sheep (Coltman, Pilkington et al. 2001, Clutton-Brock and Pemberton 2004). Gastrointestinal nematodes (GIN) are common in Soay sheep throughout life, with virtually 100 % prevalence in lambs, and immunity to GIN develops with age (Craig, Pilkington et al. 2006). GIN are a major selective force on the Soay sheep (Gulland and Fox 1992, Craig, Pilkington et al. 2006, Hayward, Wilson et al. 2011) and GIN burden, measured as FEC, is negatively associated with body weight (Coltman, Pilkington et al. 2001) and over-winter survival (Gulland and Fox 1992, Hayward, Wilson et al. 2011). Immunoglobulin isotypes IgA, IgE and IgG are involved in the acquired immune response to GIN in sheep (Stear, Strain et al. 1999, Lee, Munyard et al. 2011, Hayward 2013). Parasite-specific IgA acts at mucosal surfaces and is known to reduce worm growth and fecundity (Stear, Strain et al. 1999, Gutierrez-Gil, Perez et al. 2010, Lee, Munyard et al. 2011). Parasite-specific IgE also acts predominantly at mucosal surfaces and is involved in the degranulation of mast cells, which are white blood cells involved in parasite expulsion (McNeilly, Devaney et al. 2009, Murphy, Eckersall et al. 2010). IgG is the primary plasma antibody that can interact directly with the parasite. In Soay sheep, anti-T. *circ* IgG is positively associated with increased survival in adult females (Nussey, Watt et al. 2014, Watson, McNeilly et al. 2016, Sparks, Watt et al. 2018). However, a recent study decomposed the association between IgG and female survival into within-individual and between-individual effects and found the association was driven by within-individual variation late in life linked to senescence rather than by between-individual differences determined by genetics or early-life conditions (Froy, Sparks et al. 2019). We aim to determine whether MHC-phenotypic trait associations can indicate contemporary selection on MHC genes in Soay sheep by answering the following questions: 1) which phenotypic traits are associated with MHC class II variation? 2) Does the association between MHC variation and phenotypic traits vary with age and sex? 3) If there are associations which selection mechanism can be inferred from such associations? 4) Are there coherent patterns between MHC-phenotypic trait associations and MHC-fitness associations?

## Materials and methods

### MHC data

The genetic data used in this study was obtained from a previous study (Dicks, Pemberton et al. 2019, Dicks, Pemberton et al. 2020). Seven expressed loci (*DRB1, DQA1, DQA2, DQA2-like, DQB1, DQB2* and *DQB2-like*) within the MHC class IIa region were characterised in 118 Soay sheep using sequence-based genotyping. A total of eight MHC haplotypes were identified (named A to H) and confirmed in an additional 94 Soays selected from the pedigree to maximise diversity. A panel of 13 SNPs located in the region of MHC class IIa haplotypes, including 11 SNPs from the Ovine Infinium HD chip and two other SNPs located within the *DQA1* gene, were selected and genotyped in 5951 Soay sheep using Kompetitive Allele-specific PCR (KASP) to impute the eight haplotypes. After quality control, which included pedigree checking, the diplotypes of 5349 individuals sampled between 1985 and 2012 were identified. For each individual successfully diplotyped, the functional divergence between an individual’s two haplotypes (MHC divergence) was measured as the proportion of the amino acid sequence that differed between the two MHC haplotypes (p-distance)(Henikoff 1996, Huang, Dicks et al. 2020).

### Phenotypic traits

During an annual August catch when we try to catch as many individuals as possible, all sheep were weighed to the nearest 0.1 kg.

In our study, faecal samples were collected from as many individuals as possible at capture in August, and counts of nematode worm eggs were performed using a modified McMaster technique (MAFF 1986). This protocol enumerates Strongyle-type eggs per gram of wet weight faeces but does not differentiate between several morphologically indistinguishable species. Of the species contributing to FEC on St Kilda, *Trichostrongylus axei, Trichostrongylus vitrinus* and *Teladorsagia circumcincta* eggs are the most abundant, but eggs from *Chabertia ovina, Bunostomum trigonocephalus* and *Strongyloides papillosus* may also be present (Craig, Pilkington et al. 2006). This measure of FEC is correlated with adult worm count within the abomasum in Soay sheep on St. Kilda and on Lundy (Grenfell, Wilson et al. 1995).

IgA, IgE and IgG activity against antigens of the third larval stage of T. circ were measured using ELISA from blood samples obtained from sheep caught in the August catches. Full laboratory details and quality control measures were described in the previous study (Sparks, Watt et al. 2018). Of 3189 individuals assayed, 13 individuals failed quality control for IgA, 8 individuals for IgE and 27 individuals for IgG. Due to the lack of standard solutions, all results were measured as optical density (OD) values. The OD ratio of each sample was calculated as (sample OD – blank OD) / (positive control OD –blank OD). If the blank OD was greater than the sample OD, the OD ratio was set to zero to prevent negative OD ratios. The mean OD ratio was then calculated for each duplicated sample, and these values were used in all subsequent analyses.

For all the phenotypic traits used in this study, we only included individuals which were successfully genotyped for MHC class II diplotype, and we excluded individuals which had received an experimental treatment prior to sampling, for example, an anthelminthic bolus. The number of records that we used for each analysis is shown in Table 2.

**Table 2.** Number of records of phenotypic traits used in our study. Sample sizes for adult traits are shown as N records or N records / N individuals. Years of measurement are shown in brackets.

| Phenotypic Trait | Lambs | Yearlings | Adults/Older |
| --- | --- | --- | --- |
| **August Weight** | 1494 (1986-2012) | 841 (1985-2013) | 2599/908 (1985-2016) |
| **Strongyle FEC** | 1183 (1988-2012) | 704 (1988-2013) | 2205/794 (1988-2016) |
| **IgA** | 1404 (1990-2012) |  | 3032/1114 (1990-2015) |
| **IgE** | 1506 (1990-2012) |  | 3034 /1114 (1990-2015) |
| **IgG** | 1503(1990-2012) |  | 3018/1112(1990-2015) |

### Statistical analysis

We used animal models (AMs) to study the associations between MHC haplotypes and phenotypic traits. An AM was built for each phenotypic trait by fitting an additive genetic effect with a covariance structure proportional to the pedigree relatedness matrix in addition to the generalized linear mixed models. Each null model included fixed and random effects relevant to the studied phenotypic trait and age-group based on the findings of previous studies (Hayward, Wilson et al. 2011, Berenos, Ellis et al. 2016, Sparks, Watt et al. 2018) (Table 3). Genome-wide inbreeding, F_grm_ (Fhat3 from (Yang, Lee et al. 2011) based on 38K genome wide SNPs (Berenos et al 2016), was included in all models as a fixed effect to ensure any MHC heterozygosity associations did not simply reflect inbreeding depression. First, for each null model, we added MHC effects as MHC heterozygosity (0 for homozygote, 1 for heterozygote) and each MHC haplotype as dosage (0, 1 or 2) to test whether there are non-additive dominance effects (Hu, Deutsch et al. 2015, Lenz, Deutsch et al. 2015). In order to test any sex-dependent association between MHC diversity and phenotypic traits, we also fitted MHC by sex interactions including heterozygosity by sex and haplotype by sex interactions for each model. In all models, haplotype H was treated as a reference haplotype so any differences between individual haplotypes were relative to haplotype H. In another sets of models, we also tested the association between MHC divergence and phenotypic traits by adding MHC divergence and divergence by sex interaction into each null model.

**Table 3.** Fixed and random effects fitted in null models of each of the phenotypic traits. A ‘x’ indicates that the effect was fitted.

| Effects |  | Weight |  |  | Strongyle FEC |  |  | Immunoglobulins |  |
| --- | --- | --- | --- | --- | --- | --- | --- | --- | --- |
|  |  | Lambs | Yearlings | Adults | Lambs | Yearlings | Adults | Lambs | Older |
| Fixed effects | Sex | x | x | x | x | x | x | x | x |
|  | Litter size | x |  |  | x |  |  | x |  |
|  | Age | x |  | x | x |  | x | x | x |
|  |  | (Days) |  | (Years) | (Days) |  | (Years) | (Days) | (Years) |
|  | Age<sup>2</sup> |  |  | x |  |  | x |  | x |
|  | F<sub>grm</sub><sup>a</sup> | x | x | x | x | x | x | x | x |
|  | Maternal age | x |  |  | x |  |  | x |  |
|  | Maternal age<sup>2</sup> | x |  |  | x |  |  | x |  |
| Random effects | ID |  |  | x |  |  | x |  | x |
|  | Birth year | x |  | x | x |  | x | x | x |
|  | Capture year |  | x | x |  | x | x |  | x |
|  | Pedigree | x | x | x | x | x | x | x | x |
|  | Maternal ID | x |  |  | x | x | x | x |  |
|  | ELISA plate ID |  |  |  |  |  |  | x | x |
|  | ELISA plate run date |  |  |  |  |  |  | x | x |

<sup>a</sup> A genomic inbreeding coefficient (Yang, Lee et al. 2011),

We used a conservative statistical framework to determine the significance of MHC effects on phenotypic traits in Soay sheep. For models including MHC heterozygosity and haplotypes, the significance of MHC heterozygosity and MHC heterozygosity by sex interactions can be directly determined by whether the 95% credibility interval overlapped with zero. The significance of specific haplotypes was first determined using Wald tests for models with or without all MHC haplotypes fitted in the same model. When the Wald test was significant (p <0.05) we examined the significance of specific MHC haplotypes by conducting an additional analysis comparing the estimated effect of each haplotype against the mean of the effect estimates of all the other haplotypes (see Supplementary 1 for detailed methods). Similarly, Wald tests for each model with and without haplotype by sex interactions in the same model were used to determine the significance of haplotype by sex interactions. If the Wald test was significant, we conducted an additional analysis comparing the estimated effect of each haplotype by sex interaction with the mean of the effect estimates of all the other haplotype by sex interactions (see Supplementary 1 for detailed method). For models including only MHC divergence, the significance of MHC divergence and MHC divergence by sex interactions can be directly determined by whether the 95% credibility interval overlapped with zero. Finally, if any MHC effect by sex interaction was significant, we ran sex-specific models to determine whether there was a significant association between the MHC effect and phenotypic trait within either females or males.

Since the phenotypic traits have different distributions, we used different transformations for each trait. For August weight, we used the raw data as the distributions were normally distributed in all age groups. However, as mean August weight is known to vary with age, separate models were run for the different age classes (lambs, yearlings and adults) (Supplementary Fig 2.1) (Wilson, Pemberton et al. 2007). Strongyle FEC was not normally distributed so we carried out log transformation as Log(FEC+50) by adding half the minimum detection limit (100 eggs/g), and we again ran separate models for lambs, yearlings and adults as means were different (Supplementary Fig 2.2). Distributions of immunological traits were largely normal except lamb IgE. However, lamb IgE was not transformed in this study as MCMCglmm takes a Bayesian approach to mitigate the effect of non-Gaussian response variables. A pronounced increase in antibody levels occurs between lambs and yearlings (Sparks, Watt et al. 2018) but there was no difference in the distributions between yearlings and adults (Supplementary Fig 2.3). Thus, we fitted separate models for lambs and older sheep (including yearlings and adults) All models were run in MCMCglmm in R v.3.5.2 for 60000 iterations with a sampling interval of 30 after 5000 iterations (Hadfield 2010, R Core Team 2013).

## Results

### August weight

There was no significant association between MHC heterozygosity or divergence and August weight in any age group (Figure 1, Supplementary 3 and 4). There was no significant heterozygosity by sex interaction. However, there was an MHC divergence by sex interaction on lamb weight where the effect of MHC divergence on male lamb weight was significantly more positive than that on female lamb weight (Supplementary 3 and 4). However, MHC divergence was not significant associated with lamb weight in sex-specific models (Supplementary table 5.1).

**Figure 1.**
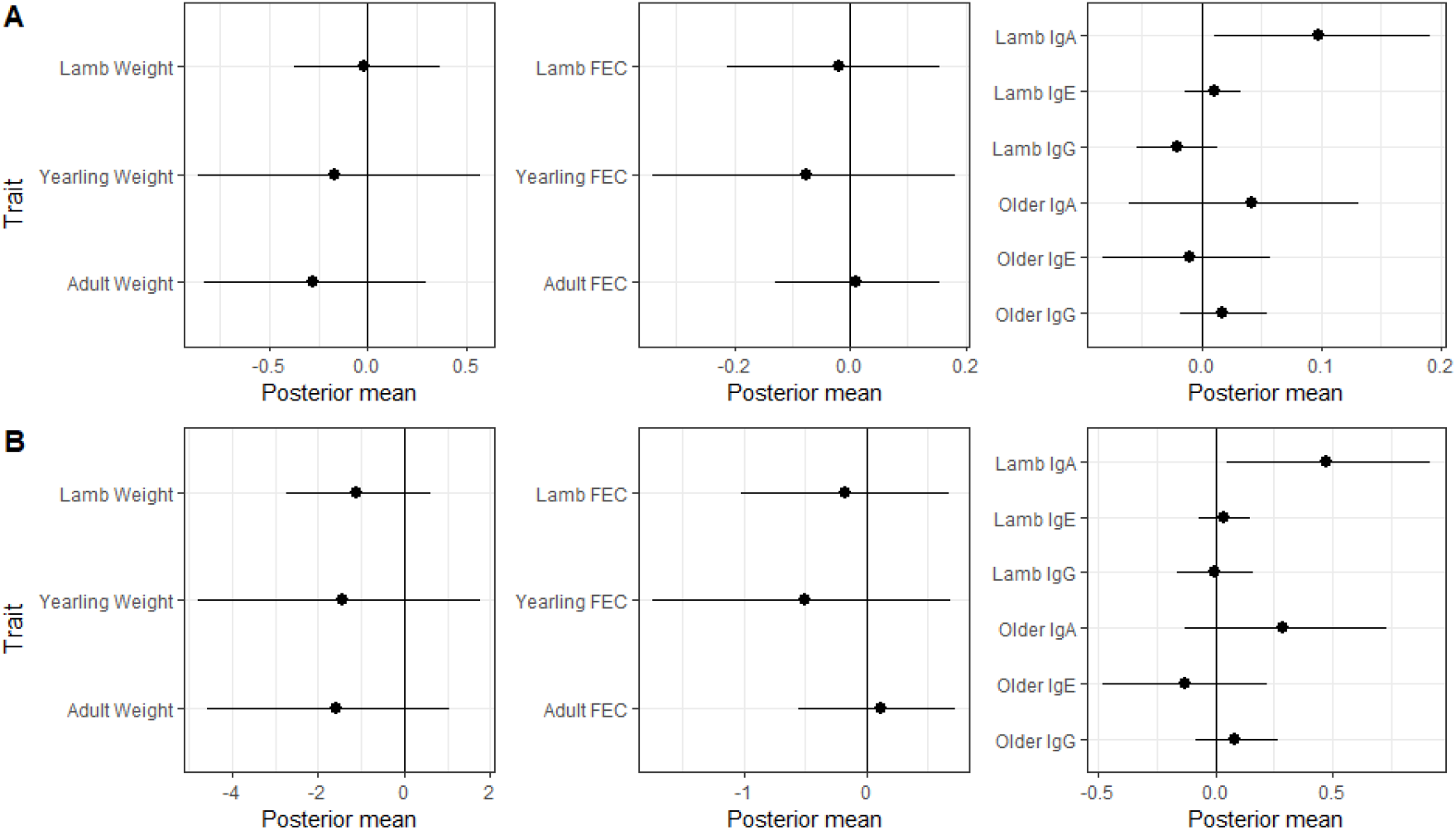
Associations between (A) MHC heterozygosity and (B) MHC divergence and phenotypic traits in Soay sheep. Results are presented as posterior means of the sampled iterations with the 95% credible intervals. Significance can be assumed if the 95% credible intervals do not overlap with zero.

The Wald test for haplotype differences was not significant in any age group so no significant association was identified between specific MHC haplotypes and August weight (Table 4, Figure 2). The result of Wald test indicated that there was no significant haplotype by sex interaction for August weight (Supplementary 6).

**Table 4.** Summary of results (p values) for Wald tests for associations between MHC haplotypes and trait values (d.f.=7). Bold values indicate significant associations (*p* <0.05). “–”indicates category was not tested.

| Phenotypic trait | Age class |  |  |
| --- | --- | --- | --- |
|  | Lambs | Yearlings | Adults/older |
| Weight | 0.32 | 0.23 | 0.93 |
| Strongyle FEC | 0.38 | 0.22 | 0.19 |
| Anti-Tc IgA | **0.0038** | - | **0.043** |
| Anti-Tc IgE | 0.26 | - | **0.00019** |
| Anti-Tc IgG | 0.088 | - | **0.0036** |

**Figure 2.**
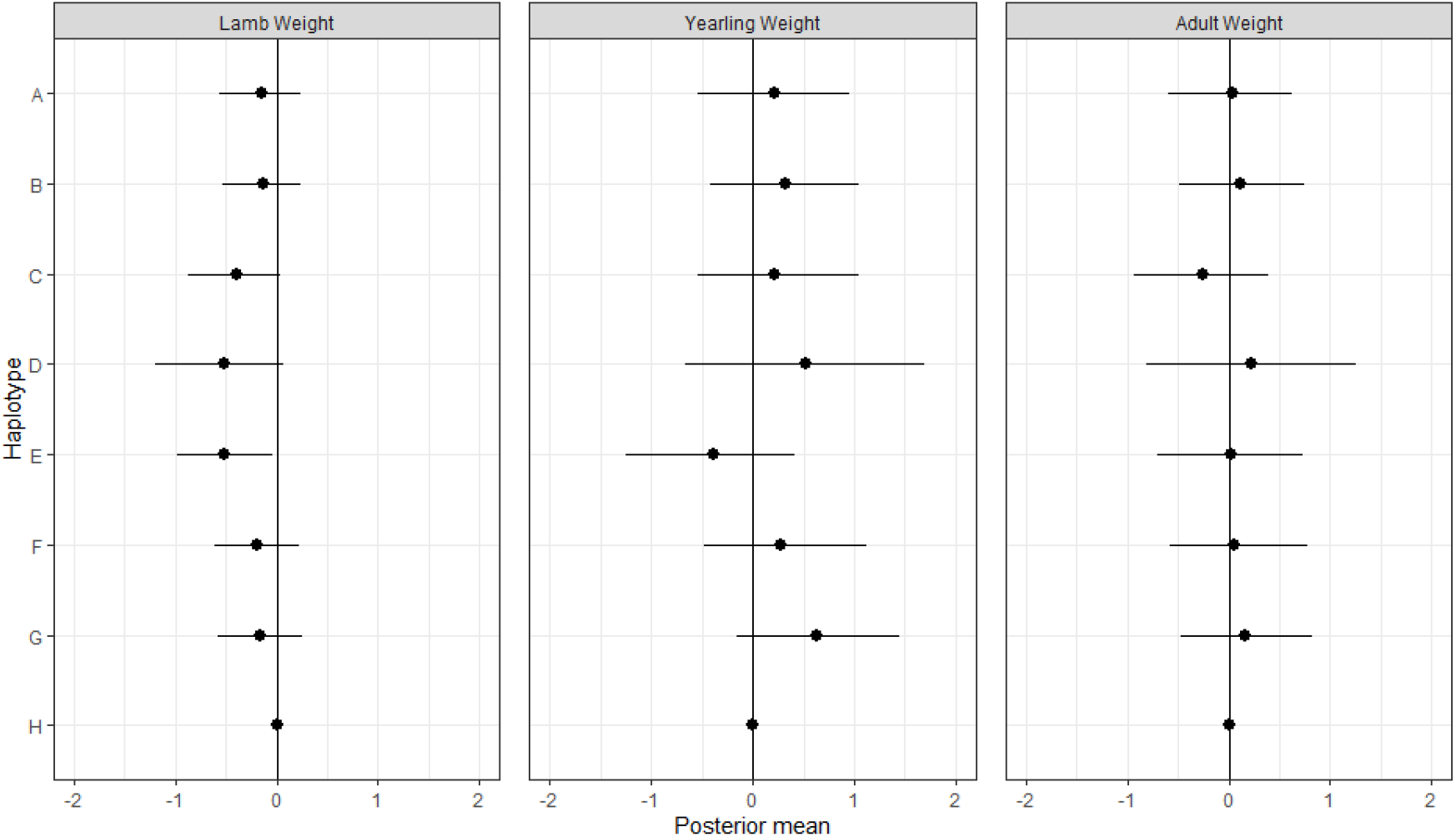
Associations between MHC haplotypes and August weight in Soay sheep. Posterior means and 95% credible intervals for each haplotype are plotted relative to haplotype H from the original model outputs. None of the Wald tests for these models were significant.

### Strongyle FEC

There was no association between MHC heterozygosity or divergence and strongyle FEC in any age group. Also, there was no significant MHC heterozygosity by sex interaction and MHC divergence by sex interaction (Figure 1, Supplementary 3 and 4). The Wald test for haplotype differences was not significant in any age group, so no significant association was identified between specific MHC haplotypes and strongyle FEC (Table 4, Figure 3). Also, the Wald test indicated that there was no significant haplotype by sex interaction for strongyle FEC (Supplementary 6).

**Figure 3.**
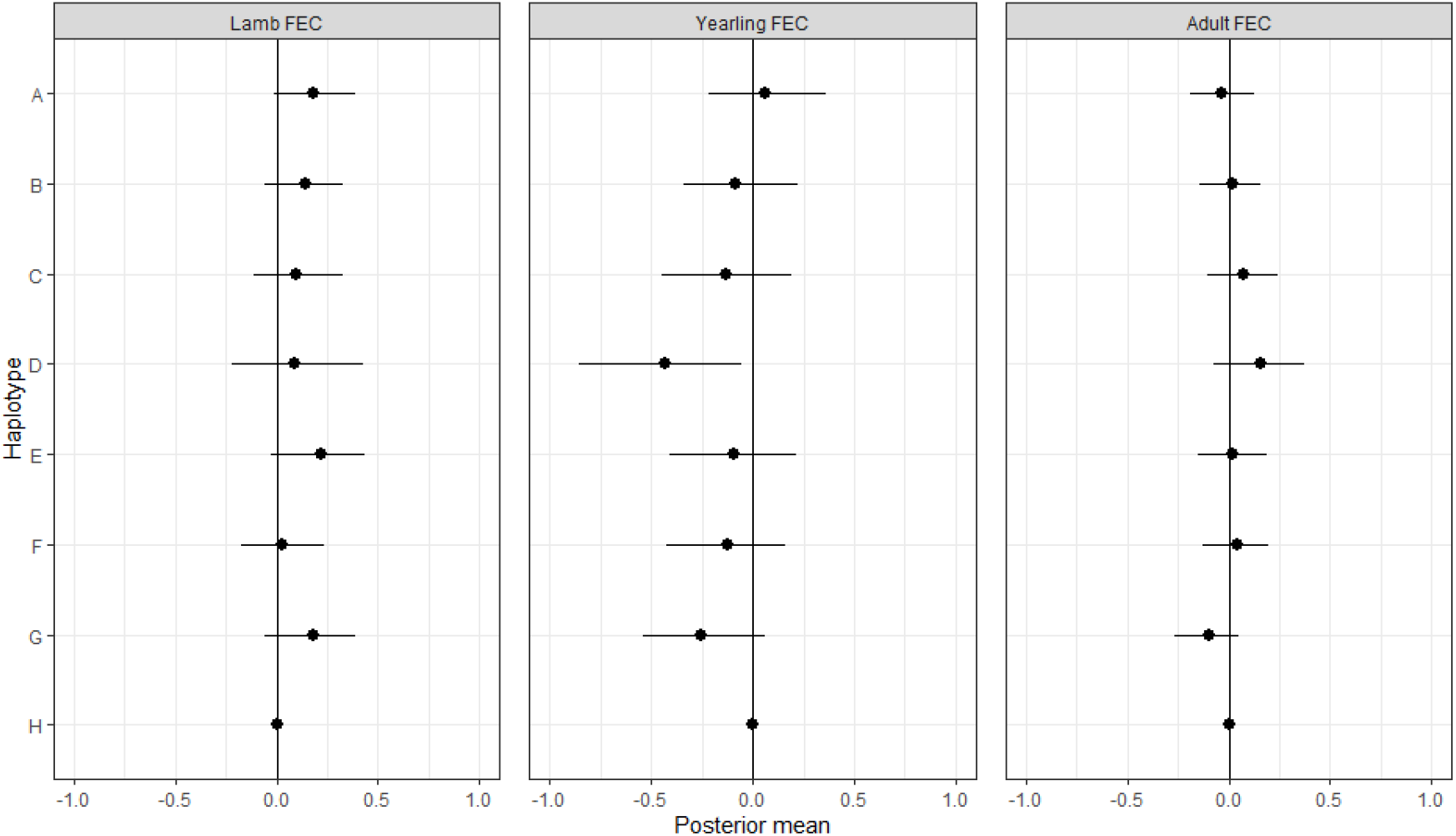
Associations between MHC haplotypes and strongyle FEC in Soay sheep. Posterior means and 95% credible intervals for each haplotype are plotted relative to haplotype H from the original model outputs. None of the Wald tests for these models were significant.

### Immunoglobulins

#### Anti-T. *circumcincta* IgA

We found that both MHC heterozygosity and MHC divergence were positively associated with IgA levels in lambs but not in older age groups (Figure 1, Supplementary 3 and 4). There was neither MHC heterozygosity by sex interaction nor MHC divergence by sex interaction (Supplementary 3 and 4).

When testing for haplotype differences, Wald tests were significant in both lambs and older sheep (Table 4). In lambs, haplotype C was associated with increased IgA while haplotype G was associated with decreased IgA (Figure 4A, Supplementary 7). In older sheep, both haplotype A was associated with decreased IgA (Figure 4B, Supplementary 5). In addition, the Wald test for haplotype*sex interactions was significant for lambs but not significant for older sheep. The effect of haplotype E on IgA in male lambs was significantly more positive than that on female lambs (Supplementary 6). When testing in sex-specific models, haplotype E was positively significant with lamb IgA level in males but not in females (Supplementary table 5.3).

**Figure 4.**
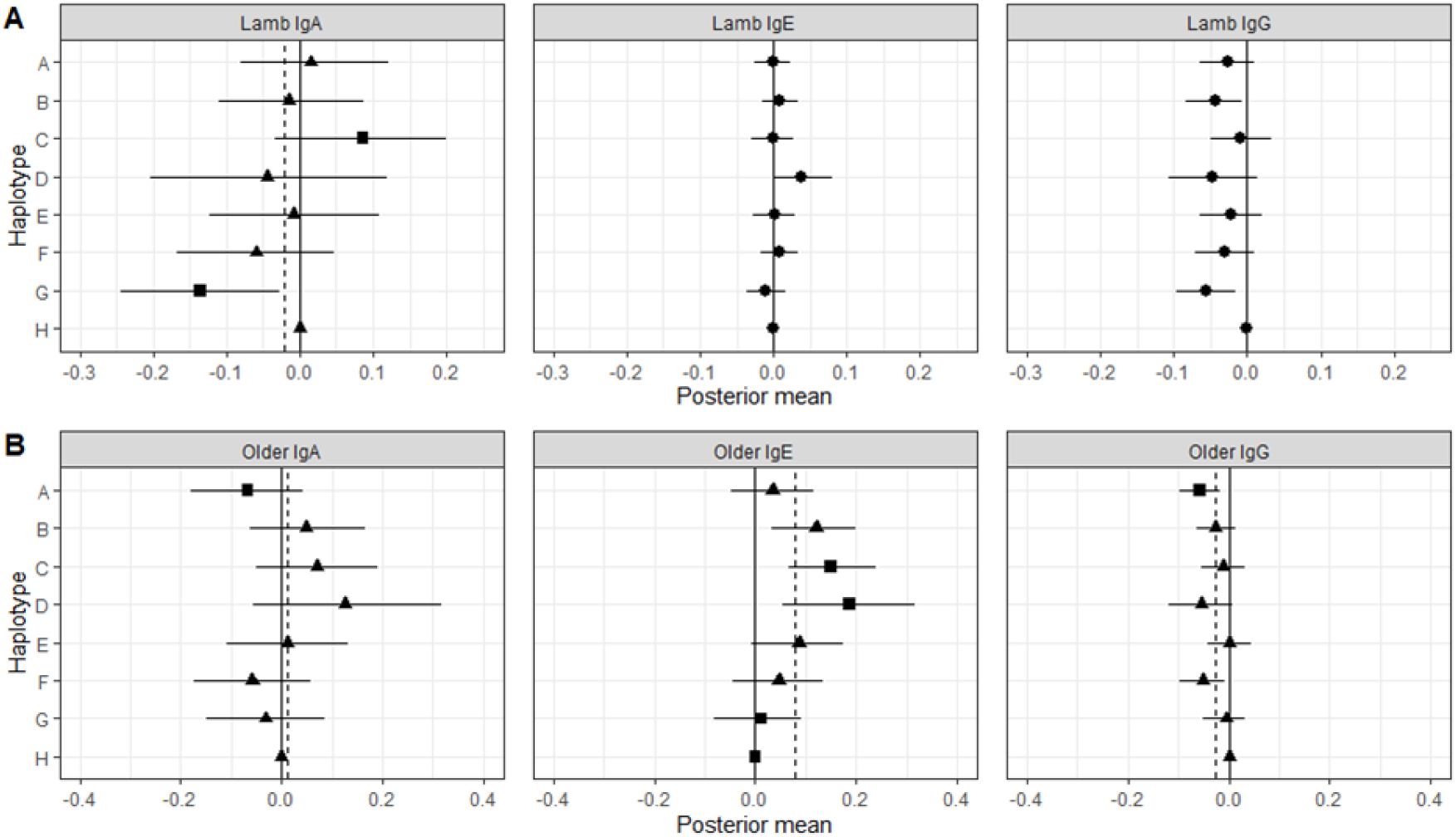
Associations between MHC haplotypes and anti-T.circ antibodies in (A) lambs and (B) older Soay sheep. Posterior means and 95% credible intervals for each haplotype are plotted relative to haplotype H from the original model outputs, with the solid vertical lines representing the haplotype H values. Circles show that Wald tests of the models were not significant, and other shapes show that Wald tests of the models were significant. When Wald tests were significant, the dashed vertical line shows the average posterior mean of all eight haplotypes, and haplotypes that differed significantly from this mean are shown as squares, while ones that did not are shown as triangles.

#### Anti-T. *circumcincta* IgE

There was no association between MHC heterozygosity or divergence and IgE levels, or significant heterozygosity by sex or divergence by sex interaction, in any age group (Figure 1, Supplementary 3 and 4).

When testing for haplotype differences, Wald tests were only significant in older sheep (Table 4). Haplotypes C and D were associated with increased IgE while haplotypes G and H were associated with decreased IgE (Figure 4B, Supplementary 7). The Wald tests for haplotype by sex interactions were not significant for either lamb or older sheep, which indicated there were no significant haplotype by sex interactions for IgE (Supplementary 6).

#### Anti-T. *circumcincta* IgG

There was no association between MHC heterozygosity or divergence and IgG in either lambs or older sheep (Figure 1, Supplementary 3 and 4). We found no significant heterozygosity by sex interaction while the effect of divergence on IgG in older sheep was significantly more negative in males than that in females (Supplementary 3 and 4). However, MHC heterozygosity was not significant associated with IgG in either older females or males in sex-specific models (Supplementary table 5.2).

When testing for haplotype differences, the Wald test was significant only in older sheep (Table 4). We found haplotype A was associated with decreased IgG (Figure 4B, Supplementary 7). However, the Wald test for haplotype by sex interactions was not significant in either lambs or older sheep, which indicated there were no significant haplotype by sex interactions for IgG (Supplementary 6).

## Discussion

In this study, we tested for associations between MHC class II variation (heterozygosity, specific haplotypes and divergence) and five representative phenotypic traits in a large sample of Soay sheep using a modelling approach that accounted for genome-wide additive genetic and inbreeding effects. While we found no associations between MHC variation and August weight or strongyle faecal egg count, we found a number of associations with anti-*T.circ*. antibody levels. Specifically, we found associations between MHC heterozygosity or divergence and divergence and IgA levels in lambs and a number of associations between specific MHC haplotypes and antibodies that vary with age, isotype and sex.

Several previous studies have found evidence that MHC heterozygosity or specific MHC alleles are associated with weight or body size in other species (e.g. (Lenz, Wells et al. 2009, Lukasch, Westerdahl et al. 2017)). Although fitness is associated with body size in Soay sheep (Coltman, Pilkington et al. 2001), we did not find any MHC associations with August weight in our study. August weight is a polygenic trait with modest heritability (Supplementary 9) (Berenos, Ellis et al. 2015). As a non-immunological trait, there is also no direct connection between the function of MHC genes and August weight. Thus, it may be not surprising that we have not found any associations between MHC variation and August weight.

MHC class II molecules are involved in the presentation of peptides from gastrointestinal nematodes for recognition by the immune system (Janeway, Travers et al. 1996). Variation in such responses to gastrointestinal nematodes infection within a population has been implicated in driving balancing selection which maintains MHC diversity (Froeschke and Sommer 2005, Madsen and Ujvari 2006, Lenz, Wells et al. 2009, Kloch, Babik et al. 2010). Association between MHC class II variation and FEC has been reported in a number of sheep breeds (reviewed in (Valilou, Rafat et al. 2015)) and previous analyses have demonstrated heritable variation in FEC as well as selection for parasite resistance in Soay sheep (Coltman, Pilkington et al. 2001, Coltman, Wilson et al. 2001, Beraldi, McRae et al. 2007, Hayward, Wilson et al. 2011). However, in this study, we did not find any association between MHC variation and strongyle FEC.

Several features of the FEC data may contribute to the lack of association in Soay sheep. The measure of FEC used here is a crude measure of parasite burden, both in terms of the way it was measured and what the measure actually represents in terms of parasite species.

Because of the dilution factor used in the modified McMasters method (MAFF 1986) for estimating FEC, the data are in multiples of 100, which combined with overdispersion, makes modelling FEC challenging. Also, individuals with 0 eggs per gram may have no strongyle worms or may have a very low burden. There may also be a very complicated relationship between FEC and the community composition of worm burden. For example in naturally infected Scottish Blackface sheep, high FEC was associated with a wider range of strongyle species (Stear, Bairden et al. 1996). Therefore, increased FEC may be confounded with increased species diversity. If there is variation among MHC haplotypes in their ability to present peptides from different worm species, we may not expect to detect a direct association between MHC haplotype and strongyle FEC.

Our results differ from an earlier study of MHC-FEC association in the same population that used the DRB1-linked microsatellite OLADRB as a marker of MHC class II haplotypes. In that study, a positive association between FEC and OLADRB allele 257 in lambs, a positive association between FEC and OLADRB allele 267 in yearlings and a negative association between FEC and OLADRB allele 263 in adults were identified (Paterson, Wilson et al. 1998). However, we did not recover such associations in the current study. The previous study used 370 individuals, whereas in this study, we used FEC measures from 1183 lambs, 704 yearlings and 2205 FEC measures from 794 adults (Table 2). Also, the MHC genotyping method used in the current study captures MHC class II composition accurately, while some OLADRB alleles correspond to multiple MHC class II haplotypes (Dicks, Pemberton et al. 2020). The advances in sample size and genotyping method are likely to contribute to this disparity in results.

Previous studies have demonstrated associations between MHC variation and antibody response in wild populations (summarized in (Gaigher, Burri et al. 2019)). Some studies found significant associations (Bonneaud, Richard et al. 2005, Charbonnel, Bryja et al. 2010, Cutrera, Zenuto et al. 2011, Gaigher, Burri et al. 2019) while others did not (Ekblom, Hasselquist et al. 2013, Cutrera, Zenuto et al. 2014). A recent study suggested that the disparity of findings in previous studies examining association between MHC variation and immunocompetence is likely caused by insufficient sample size and recommended a minimum sample size of 200 individuals to achieve sufficient power for testing associations of small effect size between MHC variation and immunocompetence (Gaigher, Burri et al. 2019). From this perspective, our sample size was large enough to have sufficient statistical power to test associations between MHC variation and immune measures (Table 2). In addition, previous studies investigating the association between MHC variation and antibody response have mainly used non-specific challenge, such as sheep red blood cell antigens (SRBC) and hemagglutinin, to elicit an antibody response (Bonneaud, Richard et al. 2005, Cutrera, Zenuto et al. 2011, Cutrera, Zenuto et al. 2014, Gaigher, Burri et al. 2019). An exception is a study of Great snipe (*Gallinago media*) that used diphtheria and tetanus toxoid as antigens (Ekblom, Hasselquist et al. 2013). Antibody against such challenges may be informative on the host’s general immunocompetence, but may not reflect the host’s immunocompetence in response to the actual pathogens imposing selection in the host’s natural environment. By using the antigen of *Teladorsagia circumcincta*, our study was able to test association between MHC variation and a relevant pathogen-specific antibody response. Although we did not find significant association between MHC variation and FEC, we still expect to find significant associations between MHC variation and anti *T.circ* antibodies because of the functional link between MHC genes and adaptive immune response (Janeway, Travers et al. 1996). Indeed, we found several associations between MHC variation and anti-*T.circ* antibodies in Soay sheep, in contrast to our results for weight and FEC. Such results suggest MHC class II variation could contribute the inter-individual heterogeneity of immune response to GIN in Soay sheep.

The associations between MHC class II variation and anti-*T.circ* antibodies varied between different age classes and among different isotypes. First, we only found association between MHC heterozygosity or divergence and IgA level in lambs. This result is consistent with a previous study of MHC-fitness associations which identified a positive association between MHC divergence and juvenile survival (Huang, Dicks et al. 2020) and indicates that there may be an age-dependent MHC heterozygote advantage or MHC divergent allele advantage in Soay sheep acting via IgA. However, the model including MHC heterozygosity, individual MHC haplotypes and MHC divergence showed neither MHC heterozygosity nor MHC divergence was significant (Supplementary 8). Thus, we could not conclude the positive effect of MHC divergence on lamb IgA was independent of MHC heterozygosity and *vice versa*. Second, although we found associations between specific MHC haplotypes and the IgA titre in both lambs and older sheep, IgE and IgG titres were only associated with MHC haplotypes in older sheep. This is consistent with a previous study in domestic sheep which found that associations with specific MHC haplotypes were only present for lamb IgA but not for lamb IgE (Ali, Murphy et al. 2019). Since the acquired immune response develops over the first year of life (Stear, Strain et al. 1999), the change in adaptive immune response may result in significant associations with specific MHC haplotypes in older IgE and IgG. Nevertheless, in terms of selection mechanism, our antibody results are consistent with negative-frequency dependent selection or fluctuating selection acting in both age classes.

The MHC-antibody associations are also partially consistent with the genetic architecture of anti-*T.circ* antibodies described in a recent genome-wide association study (GWAS) of Soay sheep. Lamb IgA and older sheep IgE levels were both associated with the MHC class II region on chromosome 20 (Sparks, Watt et al. 2019) and we recovered those associations in this study. However, the associations with IgA and IgG level in older sheep were not detected in the previous GWAS study. An explanation for such difference probably lies in the different approaches. Even with high density SNPs, a GWAS tests each SNP independently, and each SNP has only two alleles. The ovine SNP arrays have sparse coverage of the MHC and it requires ~13 SNPs to define the eight haplotypes in this region as we have found (Dicks, Pemberton et al. 2020). Therefore, it is not surprising that a GWAS could miss MHC-antibody associations in this hypervariable region.

In order to test whether there are sex-dependent associations between MHC variation and phenotypic traits, we fitted MHC by sex interactions throughout our statistical models. We found the effect of MHC divergence on lamb weight and older IgG level were significantly different between males and females. However, MHC divergence was neither associated with lamb weight nor older IgG level when testing in sex-specific models. Regarding specific MHC haplotypes, we found that only the association between haplotype E and lamb IgA was significantly different between males and females. When testing in sex-specific models, haplotype E was positively significant with lamb IgA level in males but not in females. These results suggest sex-dependent effects of haplotype E on lamb IgA level in Soay sheep which could result from the differences in ranging behaviour and life history between males and females (Clutton-Brock and Pemberton 2004).

In the present study, we have investigated MHC-phenotypic trait associations for multiple phenotypic traits. Most of the traits have been found to be associated with fitness or fitness components in previous studies (Coltman, Pilkington et al. 2001, Hayward, Wilson et al. 2011, Sparks, Watt et al. 2018). Thus, we hypothesised that we would find links between MHC-phenotypic trait associations and MHC-fitness associations. In this study, we only found associations between MHC class II genes and antibody titres. Neither August weight nor FEC was associated with MHC class II genes. When comparing MHC-antibody associations with the associations between MHC variation and fitness components identified in a previous study (Huang, Dicks et al. 2020), we found MHC divergence were both positively associated with both lamb IgA level and juvenile survival. In term of specific MHC haplotype, haplotype C was positively associated with both older IgE level but negatively associated with adult male breeding success (including both yearling and adult sheep). However, haplotype F was associated with decreased adult female life span but not associated with antibodies (Huang, Dicks et al. 2020). Although lamb IgA level was not significantly associated with juvenile survival, there is a coherent pattern between lamb IgA level and FEC (Sparks, Watt et al. 2018). A raised level of IgA is negatively associated with lamb FEC and lamb FEC is negatively associated with annual fitness (Hayward, Wilson et al. 2011, Sparks, Watt et al. 2018). Thus, it is likely that lambs with divergent MHC constitution have a survival advantage through raised anti-*T. circ* IgA level. However, such a coherent pattern is not observed in adults as older IgE level was not associated with adult fitness component (Sparks, Watt et al. 2018). Therefore, it is not clear whether selection on MHC variation could act through anti-*T.circ* antibody response.

Overall, we can conclude three points from our study. First, we only identified associations between MHC variation and immune traits, suggesting that associations are more likely to be found as one moves from highly integrative traits such as body weight to specific and more molecular traits. Second, associations between antibody traits and MHC variation varied with age, isotype and sex. Associations suggestive of divergent allele advantage and divergent allele advantage were only found in lambs, while associations suggestive of negative frequency dependent selection or fluctuating selection were found in both lambs and older sheep. Third, we found few MHC-phenotypic trait associations that were coherent with MHC-fitness associations except an association between MHC divergence and lamb IgA level. Overall, our results suggest that examining the association between MHC variation and pathogen-specific immune response is useful in the study of selection on MHC variation in wild populations.

## Acknowledgements

We thank the National Trust for Scotland for permission to work on St. Kilda and QinetiQ, Eurest and Kilda Cruises for logistics and support. We thank I. Stevenson and many volunteers who have collected samples and all those who have contributed to keeping the project going. The MHC diplotyping method and the diplotype dataset were developed and generated by Kara Dicks and Susan Johnston. SNP genotyping was conducted at the Wellcome Trust Clinical Research Facility Genetics Core. Field data collection has been supported by NERC (NE/M002896/1) over many years, the diplotyping was supported by the BBSRC and Royal Society and most of the SNP genotyping was supported by the European Research Council (AdG 250098). Wei Huang is supported by Edinburgh Global research scholarship.

## Data Accessibility

All the data and R script pf this manuscript are available through the following link: dx.doi.org/10.6084/m9.figshare.14401718

## Author contribution

W.H and J.M.P designed the study. K.L.D conducted the MHC genotyping. A.M.S and K.W generated the antibody data. J.G.P collected the data from the field. W.H analysed the data and wrote the manuscript. All the authors contributed to the final version of the manuscript.

