## Supplementary Material for "Associations between MHC class II variation and phenotypic traits in a free-living sheep population"

**Table of Contents:**

| Additional tests to verify the significance of individual haplotypes  and haplotype by sex interactions | Page 3 |
| --- | --- |
| Distribution of phenotypic traits | Page 4-6 |
| Original output of statistical models (MHC heterozygosity and  individual MHC haplotypes) | Page 7-10 |
| Original output of statistical models (MHC divergence) | Page 11,12 |
| Results of Sex-specfic models | Page 13-15 |
| Results of Additional tests to verify significance of haplotype  by sex interactions | Page 16 |
| Results of additional tests to verify significance of individual haplotypes | Page 17 |
| Results of the model including both MHC heterozygosity  and MHC divergence | Page 18 |
| Heritability of phenotypic traits | Page 19-21 |

Supplementary 1 Additional tests to verify the significance of individual haplotypes and haplotype by sex interactions

The posterior mean and covariances of the haplotype effects were used to construct a Wald test to evaluate whether trait measures differed systematically between individuals with different haplotypes. When Wald tests indicated significant differences between haplotypes, we conducted an additional test to confirm which haplotype(s) differed from the rest. We did this by obtaining the posterior distribution of the difference between the effect of a focal haplotype and the average effect of non-focal haplotypes. An MCMC p-value was calculated as *2p*, where *p* is either the posterior probability that the difference is less or greater than zero, whichever is smaller. The posterior probability is approximated by the proportion of MCMC samples that crossed zero. The same method was also used to verify the significance of haplotype by sex interactions.

Supplementary 2 Distribution of phenotypic traits

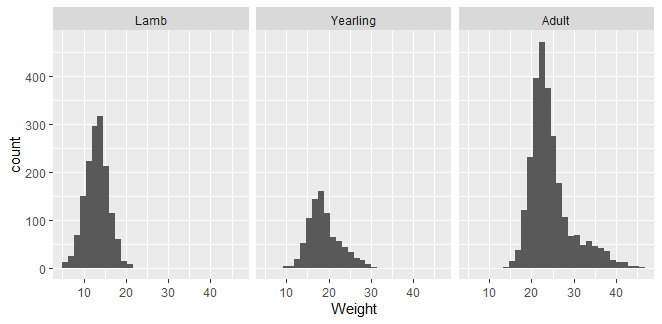

Figure 2.1. Distribution of August weight in Soay sheep.

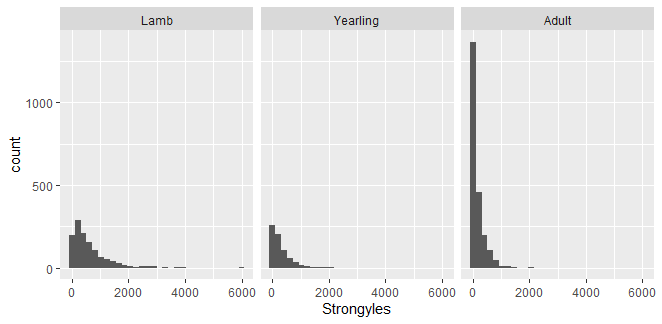

Figure 2.2. Distribution of strongyle faecal egg count (FEC) in Soay sheep.

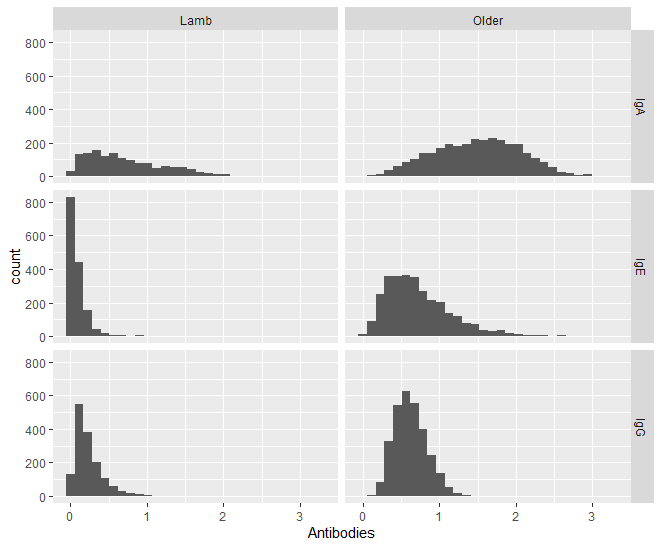

Figure 2.3. Distribution of anti-T.circ antibodies IgA, IgE and IgG in Soay sheep.

Supplementary 3 Original output of statistical models (MHC heterozygosity and individual MHC haplotypes)

Table 3.1 Results of MHC-Weight and MHC-FEC associations.

| Fixed effect | | Weight | | | FEC | | |
| --- | --- | --- | --- | --- | --- | --- | --- |
|  |  | Lamb | Yearling | Adult | Lamb | Yearling | Adult |
| Age | Estimate | 0.049 |  | 1.594 | -0.004 |  | -0.261 |
| Age | Lower 95% CI | 0.033 |  | 1.494 | -0.012 |  | -0.317 |
| Age | Upper 95% CI | 0.066 |  | 1.714 | 0.004 |  | -0.206 |
| Age^2^ | Estimate |  |  | -0.105 |  |  | 0.018 |
| Age^2^ | Lower 95% CI |  |  | -0.114 |  |  | 0.014 |
| Age^2^ | Upper 95% CI |  |  | -0.097 |  |  | 0.023 |
| Fgrm | Estimate | -3.837 | -8.383 | -7.374 | 0.462 | 2.148 | 0.831 |
| Fgrm | Lower 95% CI | -6.943 | -14.603 | -14.462 | -1.008 | -0.601 | -0.809 |
| Fgrm | Upper 95% CI | -0.822 | -0.960 | -1.548 | 2.137 | 4.665 | 2.359 |
| Matage | Estimate | 1.757 |  |  | -0.083 |  |  |
| Matage | Lower 95% CI | 1.605 |  |  | -0.160 |  |  |
| Matage | Upper 95% CI | 1.913 |  |  | -0.004 |  |  |
| Matage^2^ | Estimate | -0.138 |  |  | 0.006 |  |  |
| Matage^2^ | Lower 95% CI | -0.152 |  |  | 0.000 |  |  |
| Matage^2^ | Upper 95% CI | -0.126 |  |  | 0.013 |  |  |
| SexMale | Estimate | 0.940 | 5.102 | 9.013 | 0.732 | 0.283 | 0.887 |
| SexMale | Lower 95% CI | -0.165 | 3.051 | 6.962 | 0.221 | -0.377 | 0.321 |
| SexMale | Upper 95% CI | 1.889 | 7.077 | 11.157 | 1.222 | 1.089 | 1.427 |
| Litter size | Estimate | -3.206 |  |  | 0.376 |  |  |
| Litter size | Lower 95% CI | -3.443 |  |  | 0.246 |  |  |
| Litter size | Upper 95% CI | -2.946 |  |  | 0.493 |  |  |
| A | Estimate | -0.143 | 0.208 | 0.035 | 0.183 | 0.073 | -0.034 |
| A | Lower 95% CI | -0.571 | -0.544 | -0.592 | -0.010 | -0.238 | -0.188 |
| A | Upper 95% CI | 0.242 | 0.952 | 0.627 | 0.389 | 0.335 | 0.128 |
| A:SexMale | Estimate | 0.309 | -0.684 | -0.129 | -0.135 | 0.062 | 0.144 |
| A:SexMale | Lower 95% CI | -0.252 | -1.770 | -1.315 | -0.405 | -0.351 | -0.136 |
| A:SexMale | Upper 95% CI | 0.862 | 0.251 | 0.953 | 0.126 | 0.445 | 0.464 |
| B | Estimate | -0.135 | 0.328 | 0.118 | 0.141 | -0.073 | 0.021 |
| B | Lower 95% CI | -0.542 | -0.411 | -0.486 | -0.062 | -0.331 | -0.143 |
| B | Upper 95% CI | 0.240 | 1.051 | 0.746 | 0.327 | 0.190 | 0.156 |
| B:SexMale | Estimate | 0.056 | -0.515 | 0.254 | -0.149 | -0.064 | 0.146 |
| B:SexMale | Lower 95%CI | -0.465 | -1.451 | -0.950 | -0.386 | -0.446 | -0.149 |
| B:SexMale | Upper 95%CI | 0.583 | 0.589 | 1.390 | 0.148 | 0.302 | 0.441 |
| C | Estimate | -0.390 | 0.217 | -0.256 | 0.098 | -0.125 | 0.073 |
| C | Lower 95%CI | -0.881 | -0.534 | -0.935 | -0.113 | -0.416 | -0.101 |
| C | Upper 95%CI | 0.036 | 1.044 | 0.396 | 0.328 | 0.178 | 0.245 |
| C:SexMale | Estimate | 0.399 | -0.489 | 0.199 | -0.269 | 0.200 | -0.196 |
| C:SexMale | Lower 95%CI | -0.205 | -1.586 | -1.138 | -0.575 | -0.221 | -0.536 |
| C:SexMale | Upper 95%CI | 1.032 | 0.675 | 1.294 | 0.047 | 0.621 | 0.134 |
| D | Estimate | -0.515 | 0.525 | 0.227 | 0.087 | -0.399 | 0.159 |
| D | Lower 95%CI | -1.195 | -0.671 | -0.804 | -0.223 | -0.817 | -0.075 |
| D | Upper 95%CI | 0.068 | 1.701 | 1.250 | 0.423 | -0.003 | 0.376 |
| D:SexMale | Estimate | 0.410 | -0.576 | -0.806 | -0.177 | 0.358 | -0.018 |
| D:SexMale | Lower 95%CI | -0.477 | -2.170 | -2.586 | -0.583 | -0.244 | -0.497 |
| D:SexMale | Upper 95%CI | 1.235 | 1.175 | 0.933 | 0.217 | 1.012 | 0.420 |
| E | Estimate | -0.518 | -0.380 | 0.015 | 0.218 | -0.023 | 0.018 |
| E | Lower 95%CI | -0.989 | -1.253 | -0.701 | -0.032 | -0.316 | -0.147 |
| E | Upper 95%CI | -0.047 | 0.412 | 0.726 | 0.433 | 0.259 | 0.191 |
| E:SexMale | Estimate | 0.356 | -0.731 | 0.361 | -0.302 | -0.215 | -0.108 |
| E:SexMale | Lower 95%CI | -0.234 | -1.923 | -0.965 | -0.585 | -0.659 | -0.464 |
| E:SexMale | Upper 95%CI | 0.979 | 0.409 | 1.596 | 0.035 | 0.207 | 0.196 |
| F | Estimate | -0.190 | 0.280 | 0.049 | 0.027 | -0.086 | 0.041 |
| F | Lower 95%CI | -0.606 | -0.482 | -0.579 | -0.177 | -0.395 | -0.126 |
| F | Upper 95%CI | 0.227 | 1.126 | 0.783 | 0.233 | 0.185 | 0.200 |
| F:SexMale | Estimate | -0.037 | -1.124 | 0.787 | -0.128 | 0.054 | 0.097 |
| F:SexMale | Lower 95%CI | -0.612 | -2.275 | -0.343 | -0.390 | -0.347 | -0.222 |
| F:SexMale | Upper 95%CI | 0.575 | -0.054 | 1.982 | 0.167 | 0.455 | 0.394 |
| G | Estimate | -0.157 | 0.627 | 0.160 | 0.179 | -0.201 | -0.100 |
| G | Lower 95%CI | -0.586 | -0.149 | -0.472 | -0.061 | -0.481 | -0.267 |
| G | Upper 95%CI | 0.256 | 1.453 | 0.824 | 0.385 | 0.083 | 0.047 |
| G:SexMale | Estimate | 0.087 | -0.191 | 0.657 | -0.238 | -0.148 | 0.202 |
| G:SexMale | Lower 95%CI | -0.525 | -1.365 | -0.576 | -0.526 | -0.585 | -0.145 |
| G:SexMale | Upper 95%CI | 0.643 | 0.942 | 1.998 | 0.051 | 0.268 | 0.565 |
| Heterozygosity | Estimate | -0.015 | -0.166 | -0.279 | -0.019 | -0.077 | 0.009 |
| Heterozygosity | Lower 95%CI | -0.373 | -0.860 | -0.831 | -0.212 | -0.340 | -0.129 |
| Heterozygosity | Upper 95%CI | 0.366 | 0.569 | 0.296 | 0.156 | 0.182 | 0.156 |
| Heterozygosity:  SexMale | Estimate | 0.156 | 0.395 | 0.258 | -0.046 | 0.334 | -0.067 |
| Heterozygosity:  SexMale | Lower 95%CI | -0.359 | -0.702 | -0.885 | -0.312 | -0.032 | -0.355 |
| Heterozygosity:  SexMale | Upper 95%CI | 0.665 | 1.428 | 1.277 | 0.202 | 0.724 | 0.232 |

Table 3.2 Results of MHC-Antibody associations.

| Fixed effect | | IgA | | IgE | | IgG | |
| --- | --- | --- | --- | --- | --- | --- | --- |
|  |  | Lamb | Older | Lamb | Older | Lamb | Older |
| Age | Estimate | 0.006 | 0.027 | 0.003 | 0.098 | 0.005 | -0.042 |
| Age | Lower 95% CI | 0.002 | 0.010 | 0.002 | 0.087 | 0.004 | -0.051 |
| Age | Upper 95% CI | 0.010 | 0.043 | 0.004 | 0.111 | 0.007 | -0.034 |
| Age^2^ | Estimate |  | -0.001 |  | -0.006 |  | 0.002 |
| Age^2^ | Lower 95% CI |  | -0.002 |  | -0.007 |  | 0.002 |
| Age^2^ | Upper 95% CI |  | 0.000 |  | -0.005 |  | 0.003 |
| Fgrm | Estimate | -0.582 | 0.413 | -0.103 | -0.243 | -0.269 | -0.041 |
| Fgrm | Lower 95% CI | -1.389 | -0.615 | -0.293 | -1.026 | -0.577 | -0.431 |
| Fgrm | Upper 95% CI | 0.169 | 1.423 | 0.110 | 0.526 | 0.036 | 0.348 |
| Matage | Estimate | -0.026 |  | 0.005 |  | -0.003 |  |
| Matage | Lower 95% CI | -0.066 |  | -0.005 |  | -0.017 |  |
| Matage | Upper 95% CI | 0.012 |  | 0.015 |  | 0.013 |  |
| Matage^2^ | Estimate | 0.003 |  | >-0.001 |  | <0.001 |  |
| Matage^2^ | Lower 95% CI | 0.000 |  | -0.001 |  | -0.001 |  |
| Matage^2^ | Upper 95% CI | 0.006 |  | 0.001 |  | 0.002 |  |
| SexMale | Estimate | -0.121 | 0.049 | -0.007 | 0.044 | -0.063 | -0.102 |
| SexMale | Lower 95% CI | -0.368 | -0.218 | -0.073 | -0.153 | -0.168 | -0.214 |
| SexMale | Upper 95% CI | 0.142 | 0.344 | 0.054 | 0.278 | 0.028 | 0.007 |
| Litter size | Estimate | 0.064 |  | -0.010 |  | -0.023 |  |
| Litter size | Lower 95% CI | 0.004 |  | -0.025 |  | -0.046 |  |
| Litter size | Upper 95% CI | 0.129 |  | 0.007 |  | 0.000 |  |
| A | Estimate | 0.015 | -0.066 | -0.001 | 0.035 | -0.027 | -0.058 |
| A | Lower 95% CI | -0.081 | -0.180 | -0.025 | -0.048 | -0.064 | -0.098 |
| A | Upper 95% CI | 0.122 | 0.043 | 0.023 | 0.117 | 0.010 | -0.018 |
| A:SexMale | Estimate | 0.036 | 0.073 | -0.004 | -0.046 | 0.017 | 0.068 |
| A:SexMale | Lower 95% CI | -0.100 | -0.080 | -0.037 | -0.165 | -0.033 | 0.004 |
| A:SexMale | Upper 95% CI | 0.166 | 0.230 | 0.032 | 0.070 | 0.067 | 0.126 |
| B | Estimate | -0.015 | 0.049 | 0.009 | 0.121 | -0.044 | -0.026 |
| B | Lower 95% CI | -0.111 | -0.062 | -0.016 | 0.033 | -0.084 | -0.063 |
| B | Upper 95% CI | 0.088 | 0.165 | 0.033 | 0.199 | -0.008 | 0.014 |
| B:SexMale | Estimate | 0.038 | 0.018 | 0.007 | -0.024 | 0.026 | 0.047 |
| B:SexMale | Lower 95% CI | -0.083 | -0.127 | -0.029 | -0.136 | -0.022 | -0.013 |
| B:SexMale | Upper 95% CI | 0.182 | 0.181 | 0.037 | 0.093 | 0.082 | 0.107 |
| C | Estimate | 0.086 | 0.070 | -0.001 | 0.151 | -0.009 | -0.012 |
| C | Lower 95% CI | -0.035 | -0.051 | -0.030 | 0.066 | -0.050 | -0.055 |
| C | Upper 95% CI | 0.200 | 0.192 | 0.027 | 0.241 | 0.033 | 0.031 |
| C:SexMale | Estimate | 0.004 | -0.040 | 0.004 | -0.123 | -0.007 | 0.036 |
| C:SexMale | Lower 95% CI | -0.144 | -0.218 | -0.036 | -0.242 | -0.060 | -0.034 |
| C:SexMale | Upper 95% CI | 0.155 | 0.117 | 0.042 | 0.010 | 0.051 | 0.100 |
| D | Estimate | -0.045 | 0.127 | 0.038 | 0.187 | -0.048 | -0.055 |
| D | Lower 95% CI | -0.203 | -0.055 | 0.002 | 0.053 | -0.107 | -0.118 |
| D | Upper 95% CI | 0.119 | 0.318 | 0.080 | 0.315 | 0.014 | 0.007 |
| D:SexMale | Estimate | 0.073 | -0.155 | -0.016 | 0.018 | 0.022 | -0.044 |
| D:SexMale | Lower 95% CI | -0.125 | -0.424 | -0.069 | -0.181 | -0.056 | -0.150 |
| D:SexMale | Upper 95% CI | 0.276 | 0.097 | 0.037 | 0.225 | 0.101 | 0.063 |
| E | Estimate | -0.009 | 0.011 | 0.002 | 0.087 | -0.022 | 0.001 |
| E | Lower 95% CI | -0.123 | -0.108 | -0.028 | -0.009 | -0.065 | -0.043 |
| E | Upper 95% CI | 0.108 | 0.132 | 0.030 | 0.175 | 0.020 | 0.044 |
| E:SexMale | Estimate | 0.185 | -0.070 | 0.008 | -0.066 | 0.070 | 0.043 |
| E:SexMale | Lower 95% CI | 0.037 | -0.255 | -0.032 | -0.193 | 0.014 | -0.024 |
| E:SexMale | Upper 95% CI | 0.339 | 0.089 | 0.046 | 0.077 | 0.130 | 0.115 |
| F | Estimate | -0.059 | -0.058 | 0.009 | 0.047 | -0.030 | -0.051 |
| F | Lower 95% CI | -0.167 | -0.174 | -0.018 | -0.046 | -0.070 | -0.098 |
| F | Upper 95% CI | 0.048 | 0.058 | 0.034 | 0.134 | 0.009 | -0.009 |
| F:SexMale | Estimate | 0.123 | 0.035 | 0.017 | 0.013 | 0.031 | 0.013 |
| F:SexMale | Lower 95% CI | -0.027 | -0.115 | -0.019 | -0.107 | -0.025 | -0.054 |
| F:SexMale | Upper 95% CI | 0.252 | 0.197 | 0.054 | 0.139 | 0.081 | 0.078 |
| G | Estimate | -0.135 | -0.030 | -0.011 | 0.013 | -0.056 | -0.006 |
| G | Lower 95% CI | -0.244 | -0.147 | -0.037 | -0.082 | -0.097 | -0.052 |
| G | Upper 95% CI | -0.027 | 0.087 | 0.016 | 0.092 | -0.015 | 0.033 |
| G:SexMale | Estimate | 0.155 | 0.058 | 0.011 | -0.029 | 0.032 | 0.008 |
| G:SexMale | Lower 95% CI | 0.017 | -0.111 | -0.025 | -0.165 | -0.024 | -0.059 |
| G:SexMale | Upper 95% CI | 0.302 | 0.228 | 0.045 | 0.099 | 0.086 | 0.074 |
| Heterozygosity | Estimate | **0.098** | 0.041 | 0.011 | -0.009 | -0.020 | 0.018 |
| Heterozygosity | Lower 95% CI | **0.010** | -0.060 | -0.014 | -0.082 | -0.054 | -0.018 |
| Heterozygosity | Upper 95% CI | **0.190** | 0.131 | 0.033 | 0.058 | 0.014 | 0.055 |
| Heterozygosity:  SexMale | Estimate | -0.096 | -0.120 | -0.002 | 0.039 | -0.018 | -0.039 |
| Heterozygosity:  SexMale | Lower 95% CI | -0.226 | -0.283 | -0.036 | -0.076 | -0.063 | -0.099 |
| Heterozygosity:  SexMale | Upper 95% CI | 0.027 | 0.033 | 0.030 | 0.162 | 0.035 | 0.022 |

Supplementary 4	Original output of statistical models (MHC divergence)

Table 4.1 Results of MHC divergence-Weight and MHC divergenc-FEC associations.

| Fixed effect | | Weight | | | FEC | | |
| --- | --- | --- | --- | --- | --- | --- | --- |
|  |  | Lamb | Yearling | Adult | Lamb | Yearling | Adult |
| Age | Estimate | 0.050 |  | 1.601 | -0.004 |  | -0.262 |
| Age | Lower 95% CI | 0.034 |  | 1.482 | -0.011 |  | -0.322 |
| Age | Upper 95% CI | 0.066 |  | 1.709 | 0.004 |  | -0.209 |
| Age^2^ | Estimate |  |  | -0.106 |  |  | 0.018 |
| Age^2^ | Lower 95% CI |  |  | -0.115 |  |  | 0.014 |
| Age^2^ | Upper 95% CI |  |  | -0.098 |  |  | 0.023 |
| Divergence | Estimate | -1.117 | -1.429 | -1.597 | -0.171 | -0.510 | 0.112 |
| Divergence | Lower 95% CI | -2.739 | -4.788 | -4.568 | -1.019 | -1.739 | -0.561 |
| Divergence | Upper 95% CI | 0.622 | 1.751 | 1.057 | 0.675 | 0.689 | 0.718 |
| Fgrm | Estimate | -4.077 | -8.170 | -7.346 | 0.377 | 2.657 | 0.704 |
| Fgrm | Lower 95% CI | -7.262 | -15.417 | -13.445 | -1.217 | -0.155 | -0.863 |
| Fgrm | Upper 95% CI | -0.856 | -1.085 | -1.172 | 2.024 | 5.146 | 2.261 |
| Matage | Estimate | 1.754 |  |  | -0.090 |  |  |
| Matage | Lower 95% CI | 1.597 |  |  | -0.167 |  |  |
| Matage | Upper 95% CI | 1.903 |  |  | -0.006 |  |  |
| Matage^2^ | Estimate | -0.138 |  |  | 0.007 |  |  |
| Matage^2^ | Lower 95% CI | -0.151 |  |  | 0.000 |  |  |
| Matage^2^ | Upper 95% CI | -0.125 |  |  | 0.013 |  |  |
| SexMale | Estimate | 1.018 | 3.958 | 9.576 | 0.440 | 0.325 | 1.095 |
| SexMale | Lower 95% CI | 0.641 | 3.105 | 8.681 | 0.240 | 0.012 | 0.858 |
| SexMale | Upper 95% CI | 1.436 | 4.807 | 10.449 | 0.643 | 0.631 | 1.310 |
| SexMale: Divergence | Estimate | **2.613** | 2.648 | 1.260 | -0.631 | 1.596 | -0.894 |
| SexMale: Divergence | Lower 95% CI | **0.332** | -1.945 | -3.681 | -1.839 | -0.297 | -2.247 |
| SexMale: Divergence | Upper 95% CI | **5.067** | 7.970 | 6.661 | 0.543 | 3.377 | 0.366 |
| Litter size | Estimate | -3.209 |  |  | 0.391 |  |  |
| Litter size | Lower 95% CI | -3.468 |  |  | 0.273 |  |  |
| Litter size | Upper 95% CI | -2.948 |  |  | 0.511 |  |  |

Table 4.2 Results of MHC divergence-Antibody associations.

| Fixed effect | | IgA | | IgE | | IgG | |
| --- | --- | --- | --- | --- | --- | --- | --- |
|  |  | Lamb | Older | Lamb | Older | Lamb | Older |
| Age | Estimate | 0.007 | 0.027 | 0.003 | 0.099 | 0.006 | -0.042 |
| Age | Lower 95% CI | 0.002 | 0.011 | 0.002 | 0.087 | 0.004 | -0.051 |
| Age | Upper 95% CI | 0.011 | 0.045 | 0.004 | 0.111 | 0.007 | -0.034 |
| Age^2^ | Estimate |  | -0.001 |  | -0.006 |  | 0.002 |
| Age^2^ | Lower 95% CI |  | -0.002 |  | -0.007 |  | 0.002 |
| Age^2^ | Upper 95% CI |  | 0.001 |  | 0.005 |  | 0.003 |
| Divergence | Estimate | 0.469 | 0.29 | 0.033 | -0.131 | -0.002 | 0.085 |
| Divergence | Lower 95% CI | 0.047 | -0.128 | -0.07 | -0.477 | -0.162 | -0.083 |
| Divergence | Upper 95% CI | 0.91 | 0.728 | 0.148 | 0.218 | 0.162 | 0.265 |
| Fgrm | Estimate | -0.609 | 0.524 | -0.096 | -0.15 | -0.295 | -0.057 |
| Fgrm | Lower 95% CI | -1.418 | -0.377 | -0.305 | -0.933 | -0.572 | -0.437 |
| Fgrm | Upper 95% CI | 0.192 | 1.516 | 0.103 | 0.576 | 0.02 | 0.33 |
| Matage | Estimate | -0.023 |  | 0.005 |  | -0.002 |  |
| Matage | Lower 95% CI | -0.058 |  | -0.004 |  | -0.016 |  |
| Matage | Upper 95% CI | 0.02 |  | 0.016 |  | 0.014 |  |
| Matage^2^ | Estimate | 0.003 |  | 0 |  | 0 |  |
| Matage^2^ | Lower 95% CI | -0.001 |  | -0.001 |  | -0.001 |  |
| Matage^2^ | Upper 95% CI | 0.006 |  | 0.001 |  | 0.002 |  |
| SexMale | Estimate | 0.001 | 0.083 | 0.013 | -0.011 | -0.012 | -0.023 |
| SexMale | Lower 95% CI | -0.104 | -0.035 | -0.013 | -0.106 | -0.049 | -0.071 |
| SexMale | Upper 95% CI | 0.094 | 0.214 | 0.036 | 0.083 | 0.026 | 0.026 |
| SexMale: Divergence | Estimate | -0.318 | -0.745 | -0.067 | 0.087 | -0.109 | **-0.319** |
| SexMale: Divergence | Lower 95% CI | -0.911 | -1.512 | -0.209 | -0.44 | -0.353 | **-0.599** |
| SexMale: Divergence | Upper 95% CI | 0.257 | -0.03 | 0.092 | 0.681 | 0.108 | **-0.026** |
| Litter size | Estimate | 0.067 |  | -0.008 |  | -0.024 |  |
| Litter size | Lower 95% CI | 0.008 |  | -0.023 |  | -0.046 |  |
| Litter size | Upper 95% CI | 0.131 |  | 0.007 |  | 0.003 |  |

Supplementary 5 Results of Sex-specfic models

Table 5.1 Results of sex-specific models (divergence).

| Fixed effect | Estimate | Lower 95% CI | Upper 95% CI | Sex | Age | Trait |
| --- | --- | --- | --- | --- | --- | --- |
| Fgrm | -6.571 | -10.871 | -2.734 | Female | Lamb | Weight |
| Aged | 0.049 | 0.029 | 0.071 |  |  |  |
| Twin | -2.813 | -3.135 | -2.447 |  |  |  |
| Matage | 1.349 | 1.159 | 1.547 |  |  |  |
| Matage^2^ | -0.102 | -0.119 | -0.086 |  |  |  |
| Divergence | -0.966 | -2.619 | 0.574 |  |  |  |
| Fgrm | -2.640 | -7.805 | 2.211 | Male |  |  |
| Aged | 0.057 | 0.033 | 0.081 |  |  |  |
| Twin | -3.494 | -3.908 | -3.115 |  |  |  |
| Matage | 2.201 | 1.937 | 2.476 |  |  |  |
| Matage^2^ | -0.178 | -0.202 | -0.156 |  |  |  |
| Divergence | 1.644 | -0.251 | 3.509 |  |  |  |
| Age | -0.042 | -0.052 | -0.031 | Female | Adult | IgG |
| Age2 | 0.002 | 0.002 | 0.003 |  |  |  |
| Fgrm | -0.096 | -0.591 | 0.379 |  |  |  |
| Divergence | 0.075 | -0.088 | 0.264 |  |  |  |
| Age | -0.053 | -0.078 | -0.025 | Male |  |  |
| Age2 | 0.004 | 0.001 | 0.007 |  |  |  |
| Fgrm | -0.267 | -0.998 | 0.391 |  |  |  |
| Divergence | -0.264 | -0.516 | 0.005 |  |  |  |

Table 5.2 Results of sex-specific models (heterozygosity and individual haplotypes).

| Fixed effect | Estimate | Lower 95% CI | Upper 95% CI | Sex | Age | Trait |
| --- | --- | --- | --- | --- | --- | --- |
| Heterozygosity | 0.083 | -0.014 | 0.183 | Female | Lamb | IgA |
| A | 0.003 | -0.107 | 0.114 |  |  |  |
| B | -0.017 | -0.124 | 0.092 |  |  |  |
| C | 0.074 | -0.047 | 0.193 |  |  |  |
| D | -0.044 | -0.212 | 0.136 |  |  |  |
| E | -0.045 | -0.167 | 0.091 |  |  |  |
| F | -0.068 | -0.187 | 0.050 |  |  |  |
| G | -0.126 | -0.247 | -0.008 |  |  |  |
| Aged | 0.007 | 0.001 | 0.013 |  |  |  |
| Fgrm | -1.073 | -2.310 | -0.001 |  |  |  |
| Twin | 0.096 | 0.007 | 0.189 |  |  |  |
| Matage | -0.034 | -0.085 | 0.018 |  |  |  |
| Matage^2^ | 0.004 | -0.001 | 0.008 |  |  |  |
| Heterozygosity | 0.010 | -0.078 | 0.097 | Male |  |  |
| A | 0.058 | -0.046 | 0.164 |  |  |  |
| B | 0.041 | -0.063 | 0.146 |  |  |  |
| C | 0.113 | -0.008 | 0.235 |  |  |  |
| D | 0.025 | -0.130 | 0.192 |  |  |  |
| E | 0.216 | 0.091 | 0.328 |  |  |  |
| F | 0.094 | -0.019 | 0.202 |  |  |  |
| G | 0.041 | -0.076 | 0.143 |  |  |  |
| Aged | 0.007 | 0.001 | 0.012 |  |  |  |
| Fgrm | -0.181 | -1.287 | 0.960 |  |  |  |
| Twin | 0.051 | -0.030 | 0.136 |  |  |  |
| Matage | -0.004 | -0.062 | 0.065 |  |  |  |
| Matage^2^ | 0.001 | -0.005 | 0.007 |  |  |  |

| ho | 0.11192 | -0.02062 | 0.240409 | lb_iga | female |
| --- | --- | --- | --- | --- | --- |
| ha | 0.01408 | -0.10438 | 0.134368 | lb_iga | female |
| hb | -0.01426 | -0.14274 | 0.108 | lb_iga | female |
| hc | 0.076142 | -0.04699 | 0.216601 | lb_iga | female |
| hd | -0.0443 | -0.24038 | 0.140875 | lb_iga | female |
| he | -0.046 | -0.18167 | 0.087944 | lb_iga | female |
| hf | -0.06752 | -0.17944 | 0.077635 | lb_iga | female |
| hg | -0.14317 | -0.26514 | -0.00933 | lb_iga | female |
| aged | 0.007056 | 0.000622 | 0.012892 | lb_iga | female |
| Fhat3 | -1.09067 | -2.21409 | 0.07634 | lb_iga | female |
| Twin | 0.092815 | 0.001933 | 0.182147 | lb_iga | female |
| matage | -0.03323 | -0.08563 | 0.017712 | lb_iga | female |
| matage2 | 0.003647 | -0.00041 | 0.00837 | lb_iga | female |
| ho | -0.07176 | -0.20627 | 0.051303 | lb_iga | male |
| ha | 0.091809 | -0.01262 | 0.220236 | lb_iga | male |
| hb | 0.044298 | -0.07421 | 0.16172 | lb_iga | male |
| hc | 0.135077 | -0.00695 | 0.252139 | lb_iga | male |
| hd | 0.043508 | -0.14315 | 0.209012 | lb_iga | male |
| he | 0.241261 | 0.125936 | 0.380554 | lb_iga | male |
| hf | 0.104284 | -0.01992 | 0.228448 | lb_iga | male |
| hg | 0.043916 | -0.08249 | 0.165511 | lb_iga | male |
| aged | 0.006568 | 0.001075 | 0.011925 | lb_iga | male |
| Fhat3 | -0.20604 | -1.33606 | 0.923735 | lb_iga | male |
| Twin | 0.049144 | -0.03046 | 0.134918 | lb_iga | male |
| matage | -0.00452 | -0.07281 | 0.05324 | lb_iga | male |
| matage2 | 0.001037 | -0.0041 | 0.006649 | lb_iga | male |
| ho | 0.054119 | 0.000478 | 0.101577 | ad_igg | female |
| ha | -0.07829 | -0.12456 | -0.03533 | ad_igg | female |
| hb | -0.03189 | -0.07432 | 0.015337 | ad_igg | female |
| hc | -0.02126 | -0.06754 | 0.02749 | ad_igg | female |
| hd | -0.05687 | -0.1259 | 0.008529 | ad_igg | female |
| he | -0.00251 | -0.04914 | 0.043144 | ad_igg | female |
| hf | -0.06494 | -0.11076 | -0.01667 | ad_igg | female |
| hg | -0.0168 | -0.06362 | 0.028078 | ad_igg | female |
| age | -0.04208 | -0.05145 | -0.03186 | ad_igg | female |
| age2 | 0.00236 | 0.001609 | 0.003183 | ad_igg | female |
| Fhat3 | 0.083638 | -0.37266 | 0.554407 | ad_igg | female |
| ho | -0.02675 | -0.09551 | 0.038429 | ad_igg | male |
| ha | 0.021828 | -0.03846 | 0.081643 | ad_igg | male |
| hb | 0.03012 | -0.03517 | 0.08532 | ad_igg | male |
| hc | 0.029748 | -0.03117 | 0.100371 | ad_igg | male |
| hd | -0.11308 | -0.20891 | -0.02124 | ad_igg | male |
| he | 0.028126 | -0.03773 | 0.092122 | ad_igg | male |
| hf | -0.05671 | -0.11908 | 0.001064 | ad_igg | male |
| hg | -0.00145 | -0.06623 | 0.063098 | ad_igg | male |
| age | -0.05239 | -0.07884 | -0.0246 | ad_igg | male |
| age2 | 0.004082 | 0.000491 | 0.007108 | ad_igg | male |
| Fhat3 | -0.23041 | -0.97687 | 0.436418 | ad_igg | male |

Table 5.3 Results of additional tests that verify the significance of individual haplotype for sex-specific models (lamb IgA). The results of Wald tests are shown in brackets and significant haplotypes are presented in bold font.

| Haplotype | Estimate | Mean | *P* | Sex |
| --- | --- | --- | --- | --- |
| A | 0.003 | -0.028 | 0.344 | Female  (0.028) |
| B | -0.017 |  | 0.718 |  |
| **C** | **0.074** |  | **0.016** |  |
| D | -0.044 |  | 0.815 |  |
| E | -0.045 |  | 0.685 |  |
| F | -0.068 |  | 0.256 |  |
| **G** | **-0.126** |  | **0.008** |  |
| H | 0.000 |  | 0.543 |  |
| A | 0.058 | 0.073 | 0.595 | Male  (0.017) |
| B | 0.041 |  | 0.262 |  |
| C | 0.113 |  | 0.330 |  |
| D | 0.025 |  | 0.438 |  |
| **E** | **0.216** |  | **< 0.001** |  |
| F | 0.094 |  | 0.508 |  |
| G | 0.041 |  | 0.350 |  |
| H | 0.000 |  | 0.079 |  |

Supplementary 6 Results of Additional tests to verify significance of haplotype by sex interactions

Table 6.1. Summary of results (*p value*) of Wald tests for haplotype by sex interactions (d.f.=7) . Bold number indicate significant results (*p* <0.05).

| Trait | Lambs | Yearlings | Adults/older |
| --- | --- | --- | --- |
| Weight | 0.39 | 0.61 | 0.54 |
| FEC | 0.6 | 0.3 | 0.082 |
| IgA | **0.042** | - | 0.39 |
| IgE | 0.83 | - | 0.45 |
| IgG | 0.23 | - | 0.14 |

Table 6.2 Results of additional tests that verify the significance of individual haplotypeby sex interactions. Only models where Wald tests for haplotype by sex interactions were significant are shown, with the mean of estimates of all eight haplotypes for each model. Significant haplotype by sex interactions are presented in bold font.

| Trait | Haplotype | Estimate | Mean | *P* |
| --- | --- | --- | --- | --- |
| Lamb IgA | A: SexMale | 0.036 | 0.077 | 0.287 |
|  | B: SexMale | 0.038 |  | 0.264 |
|  | C: SexMale | 0.004 |  | 0.137 |
|  | D: SexMale | 0.073 |  | 0.950 |
|  | **E: SexMale** | **0.185** |  | **0.025** |
|  | F: SexMale | 0.123 |  | 0.260 |
|  | G: SexMale | 0.155 |  | 0.065 |
|  | H: SexMale | 0.000 |  | 0.152 |

Supplementary 7 Results of additional tests to verify significance of individual haplotypes

Table 7.1 Results of additional tests that verify the significance of individual haplotypes in models with pedigree fitted (AMs). Only models where Wald tests for haplotype effects were significant are shown, with the mean of estimates of all eight haplotypes for each model. Significant haplotypes are presented in bold font.

| Trait | Haplotype | Estimate | Mean | *P* |
| --- | --- | --- | --- | --- |
| Lamb IgA | A | 0.015 | -0.020 | 0.244 |
|  | B | -0.015 |  | 0.854 |
|  | **C** | **0.086** |  | **0.002** |
|  | D | -0.045 |  | 0.696 |
|  | E | -0.009 |  | 0.782 |
|  | F | -0.059 |  | 0.229 |
|  | **G** | **-0.135** |  | **<0.001** |
|  | H | 0 |  | 0.617 |
| Older IgA | **A** | **-0.066** | 0.013 | **0.023** |
|  | B | 0.049 |  | 0.264 |
|  | C | 0.070 |  | 0.147 |
|  | D | 0.127 |  | 0.116 |
|  | E | 0.011 |  | 0.959 |
|  | F | -0.058 |  | 0.053 |
|  | G | -0.030 |  | 0.260 |
|  | H | 0 |  | 0.795 |
| Older IgE | A | 0.035 | 0.080 | 0.067 |
|  | B | 0.121 |  | 0.069 |
|  | **C** | **0.151** |  | **0.013** |
|  | **D** | **0.187** |  | **0.039** |
|  | E | 0.087 |  | 0.820 |
|  | F | 0.047 |  | 0.231 |
|  | **G** | **0.013** |  | **0.017** |
|  | **H** | **0** |  | **0.020** |
| Older IgG | **A** | **-0.058** | -0.026 | **0.008** |
|  | B | -0.026 |  | 0.972 |
|  | C | -0.012 |  | 0.328 |
|  | D | -0.055 |  | 0.229 |
|  | E | 0.001 |  | 0.055 |
|  | F | -0.051 |  | 0.056 |
|  | G | -0.006 |  | 0.119 |
|  | H | 0 |  | 0.109 |
|  | G | -0.011 |  | 0.236 |
|  | H | 0 |  | 0.123 |

Supplementary 8 Results of the model including both MHC heterozygosity and MHC divergence

Table 8.1 Results of the model examining association between MHC and lamb IgA including both MHC heterozygosity and MHC divergence.

| Fixed effect | Estimate | Lower 95% CI | Upper 95% CI |
| --- | --- | --- | --- |
| A | 0.016 | -0.096 | 0.115 |
| B | -0.010 | -0.112 | 0.093 |
| C | 0.083 | -0.048 | 0.196 |
| D | -0.050 | -0.205 | 0.113 |
| E | -0.014 | -0.147 | 0.102 |
| F | -0.058 | -0.166 | 0.052 |
| G | -0.136 | -0.248 | -0.029 |
| SexMale | -0.124 | -0.367 | 0.138 |
| Aged | 0.006 | 0.002 | 0.011 |
| Fgrm | -0.596 | -1.384 | 0.251 |
| Twin | 0.066 | 0.004 | 0.130 |
| Matage | -0.026 | -0.065 | 0.013 |
| Matage2 | 0.003 | 0.000 | 0.006 |
| Divergence | 0.201 | -0.718 | 1.121 |
| Heterozygosity | 0.065 | -0.115 | 0.240 |
| SexMale:divergence | -0.470 | -1.816 | 0.821 |
| SexMale:heterozygosity | -0.018 | -0.261 | 0.258 |
| SexMale:A | 0.040 | -0.098 | 0.176 |
| SexMale:B | 0.034 | -0.111 | 0.160 |
| SexMale:C | 0.016 | -0.145 | 0.168 |
| SexMale:D | 0.091 | -0.128 | 0.288 |
| SexMale:E | 0.195 | 0.040 | 0.351 |
| SexMale:F | 0.129 | -0.019 | 0.269 |
| SexMale:G | 0.160 | 0.019 | 0.303 |

Supplementary 9	Heritability of phenotypic traits.

Table 9.1 Estimates of variance components and heritability (on the latent scale and shown in bold font) of August weight (h^2^). The variance components differed between different age classes. We show additive genetic effect (V_A_), birth year effect (V_BY_), measurement year effect (V_CY_), mother identity effect (V_M_), permanent environment effect (V_PE_), and the residual variance (V_R_).

| Trait | Component | Lower 95% CI | Upper 95% CI | Mean |
| --- | --- | --- | --- | --- |
| Lamb Weight | V_A_ | 0.395 | 1.354 | 0.817 |
|  | V_BY_ | 0.884 | 3.236 | 1.960 |
|  | V_M_ | 0.412 | 1.001 | 0.701 |
|  | V_R_ | 1.750 | 2.491 | 2.138 |
|  | **h^2^** | **0.064** | **0.238** | **0.147** |
| Yearling Weight | V_A_ | 0.457 | 3.011 | 1.666 |
|  | V_CY_ | 0.910 | 3.530 | 2.023 |
|  | V_R_ | 4.280 | 6.605 | 5.423 |
|  | **h^2^** | **0.046** | **0.317** | **0.183** |
| Adult Weight | V_A_ | 1.322 | 3.229 | 2.248 |
|  | V_BY_ | <0.001 | 0.453 | 0.197 |
|  | V_CY_ | 0.645 | 2.483 | 1.540 |
|  | V_PE_ | 2.939 | 4.622 | 3.783 |
|  | V_R_ | 2.061 | 2.339 | 2.195 |
|  | **h^2^** | **0.134** | **0.312** | **0.225** |

Table 9.2 Estimates of variance components and heritability (on the latent scale and shown in bold font) of FEC (h^2^). The variance components differed between different age classes. We show additive genetic effect (V_A_), birth year effect (V_BY_), measurement year effect (V_CY_), mother identity effect (V_M_), permanent environment effect (V_PE_), and the residual variance (V_R_).

| Trait | Component | Lower 95% CI | Upper 95% CI | Mean |
| --- | --- | --- | --- | --- |
| Lamb FEC | V_A_ | 0.045 | 0.198 | 0.119 |
|  | V_BY_ | 0.120 | 0.415 | 0.247 |
|  | V_M_ | <0.001 | 0.035 | 0.01 |
|  | V_R_ | 0.470 | 0.623 | 0.542 |
|  | **h^2^** | **0.050** | **0.217** | **0.131** |
| Yearling FEC | V_A_ | 0.061 | 0.311 | 0.178 |
|  | V_CY_ | 0.023 | 0.224 | 0.107 |
|  | V_R_ | 0.490 | 0.74 | 0.604 |
|  | **h^2^** | **0.065** | **0.342** | **0.201** |
| Adult FEC | V_A_ | 0.017 | 0.111 | 0.063 |
|  | V_BY_ | <0.001 | 0.038 | 0.018 |
|  | V_CY_ | 0.007 | 0.043 | 0.022 |
|  | V_PE_ | <0.001 | 0.087 | 0.045 |
|  | V_R_ | 0.560 | 0.645 | 0.603 |
|  | **h^2^** | **0.023** | **0.144** | **0.084** |

Table 9.3 Estimates of variance components and heritability (on the latent scale and shown in bond font) of Antibodies (h^2^). The variance components differed between different age classes. We show additive genetic effect (V_A_), birth year effect (V_BY_), measurement year effect (V_CY_), mother identity effect (V_M_), permanent environment effect (V_PE_), and the residual variance (V_R_).

| Trait | Component | Lower 95% CI | Upper 95% CI | Mean |
| --- | --- | --- | --- | --- |
| Lamb IgA | V_A_ | 0.074 | 0.139 | 0.106 |
|  | V_BY_ | 0.002 | 0.029 | 0.013 |
|  | V_M_ | 0.001 | 0.027 | 0.015 |
|  | V_PL_ | <0.001 | 0.015 | 0.007 |
|  | V_R_ | 0.075 | 0.123 | 0.098 |
|  | **h^2^** | **0.320** | **0.557** | **0.443** |
| Lamb IgE | V_A_ | 0.001 | 0.004 | 0.002 |
|  | V_BY_ | <0.001 | 0.001 | 0.001 |
|  | V_M_ | <0.001 | 0.001 | 0.001 |
|  | V_PL_ | <0.001 | 0.001 | <0.001 |
|  | V_R_ | 0.009 | 0.012 | 0.011 |
|  | **h^2^** | **0.069** | **0.241** | **0.152** |
| Lamb IgG | V_A_ | 0.005 | 0.011 | 0.008 |
|  | V_BY_ | 0.001 | 0.007 | 0.003 |
|  | V_M_ | <0.001 | 0.002 | 0.001 |
|  | V_PL_ | <0.001 | 0.003 | 0.001 |
|  | V_R_ | 0.019 | 0.025 | 0.022 |
|  | **h^2^** | **0.160** | **0.324** | **0.238** |
| Older IgA | V_A_ | 0.140 | 0.226 | 0.184 |
|  | V_BY_ | <0.001 | 0.012 | 0.005 |
|  | V_CY_ | <0.001 | 0.005 | 0.002 |
|  | V_PE_ | 0.029 | 0.076 | 0.052 |
|  | V_PL_ | 0.004 | 0.012 | 0.008 |
|  | V_R_ | 0.055 | 0.062 | 0.059 |
|  | **h^2^** | **0.500** | **0.686** | **0.593** |
| Older IgE | V_A_ | 0.052 | 0.092 | 0.071 |
|  | V_BY_ | <0.001 | 0.002 | <0.001 |
|  | V_CY_ | 0.001 | 0.006 | 0.003 |
|  | V_PE_ | 0.033 | 0.060 | 0.046 |
|  | V_PL_ | <0.001 | 0.003 | 0.001 |
|  | V_R_ | 0.036 | 0.041 | 0.038 |
|  | **h^2^** | **0.330** | **0.532** | **0.440** |
| Older IgG | V_A_ | 0.003 | 0.010 | 0.007 |
|  | V_BY_ | <0.001 | 0.001 | <0.001 |
|  | V_CY_ | <0.001 | 0.004 | 0.002 |
|  | V_PE_ | 0.013 | 0.020 | 0.016 |
|  | V_PL_ | 0.004 | 0.008 | 0.006 |
|  | V_R_ | 0.014 | 0.016 | 0.015 |
|  | **h^2^** | **0.065** | **0.221** | **0.147** |
